# Diet and temperature interactively impact brown adipose tissue gene regulation controlled by DNA methylation

**DOI:** 10.1101/2025.08.25.672110

**Authors:** Tobias Hagemann, Anne Hoffmann, Kerstin Rohde-Zimmermann, Helen Broghammer, Lucas Massier, Peter Kovacs, Michael Stumvoll, Matthias Blüher, John T. Heiker, Juliane Weiner

**Author notes:** Corresponding authors: Juliane Weiner, ^2^Leipzig University Medical Center, Divisions of Endocrinology and Nephrology, University of Leipzig, Leipzig, Germany, Liebigstr. 21, D-04103 Leipzig, Germany, John T. Heiker, Helmholtz Institute for Metabolic, Obesity and Vascular Research (HI-MAG), Helmholtz Zentrum München at the University of Leipzig and University Hospital Leipzig Philipp-Rosenthal-Straße 27, D-04103 Leipzig, Germany. These authors contribute equally (Tobias Hagemann, Anne Hoffmann, Kerstin Rohde-Zimmermann, John T. Heiker, Juliane Weiner).

## Abstract

Controlling brown adipose tissue (BAT) plasticity in adulthood holds promising potential for effective new obesity therapies by targeting the mechanisms of adaptive thermogenesis. Recent studies have shown that BAT development and function are under epigenetic control, with DNA methylation linked to the regulation of key thermogenic and metabolic genes. Here we sought to understand how diet and cold exposure interactively shape BAT gene regulation controlled by DNA methylation. Mice (N = 8 per group) were housed under cold exposure (8°C) or thermoneutrality (30°C) and fed either chow or high-fat diet (HFD). BAT was isolated for transcriptome (RNAseq) and methylome (RRBS) analyses. We identified differentially methylated and expressed genes (DMEGs) by comparing the effects of cold exposure under chow and high-fat diet, as well as by analyzing the interaction between temperature and diet. Functional pathway enrichment and EpiFactors Database screening were used to assess epigenetic regulators, and candidate gene expression was validated by modulating DNA-methylation in vitro.

We identified ∼1,360 differentially expressed genes (DEGs) uniquely affected by the diet-temperature interaction with most downregulated in HFD-fed mice, indicating that obesity limits the transcriptional response of BAT to cold. 65 DMEGs (4% of DEGs) were largely diet-specific in response to cold exposure, suggesting that DNA methylation contributes to a selective layer of gene regulation during BAT adaptation to distinct metabolic states. In HFD-fed mice, DMEGs were enriched in pathways related to mitochondrial dysfunction, altered lipid metabolism, neuroendocrine signaling, and compensatory stress responses, contrasting the adaptive thermogenic profile observed in chow-fed controls. Differentially expressed genes of epigenetic regulators such as *Tet2*, *Dnmt3a* and *Apobec1* showed diet- and cold-dependent regulation, indicating impaired methylation flexibility under obesogenic conditions. Using cell culture experiments, we confirmed the regulation of gene expression of candidate genes, validating the functional link between DNA methylation and thermogenic gene regulation. This is the first study to demonstrate an epigenetic response to cold exposure that differs by dietary condition, particularly in obesity. Our findings highlight a coordinated transcriptional and epigenetic remodeling of BAT, shaped by both environmental and metabolic signals. These insights may inform targeted epigenetic or nutritional strategies to restore BAT function and would further strengthen our understanding of how these might be used as therapeutic basis to improve metabolic health in obesity.

**Graphical abstract:** 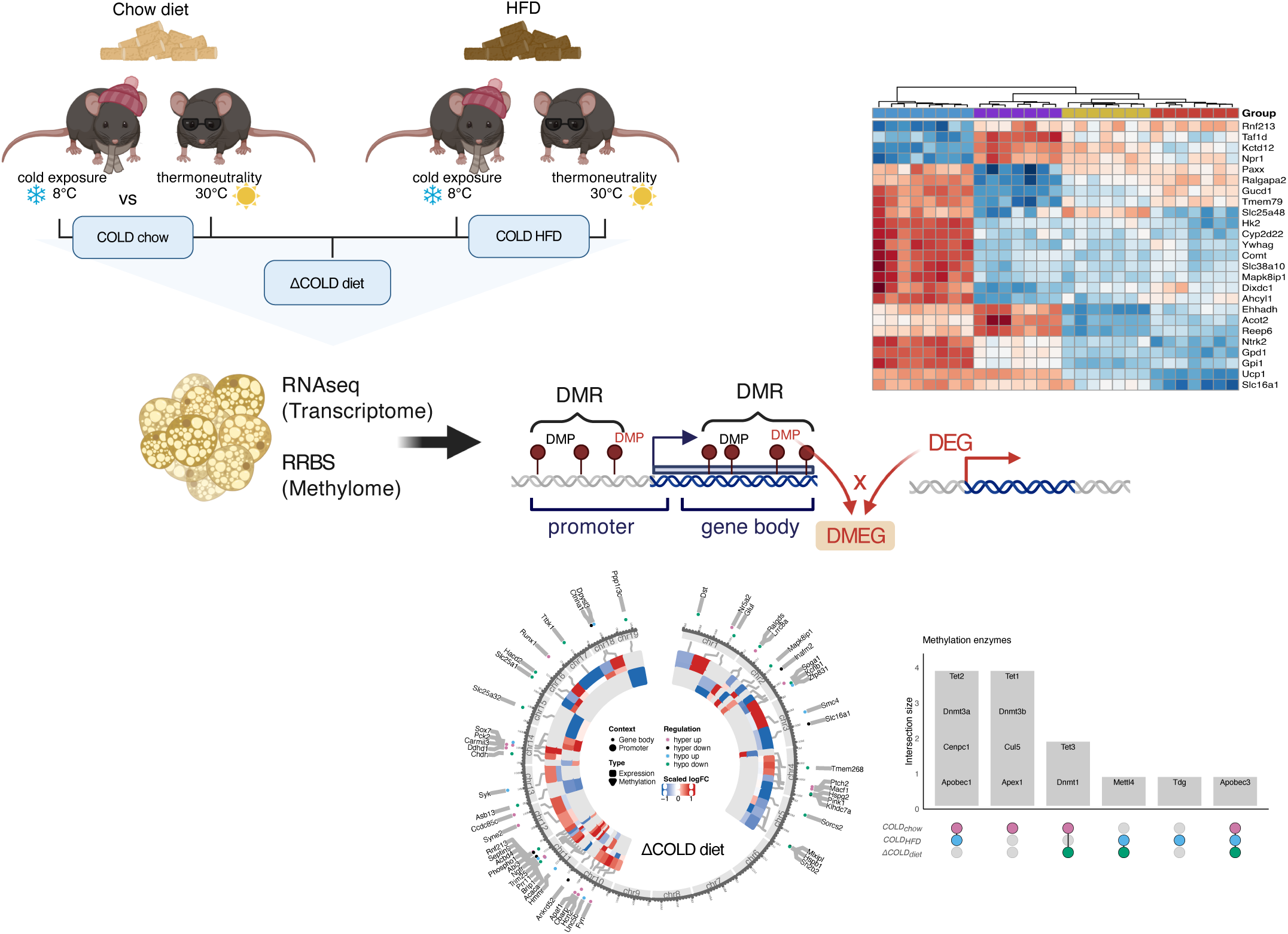

Mice were housed under cold exposure or thermoneutrality and fed either chow or high-fat diet (HFD). RNA sequencing (RNAseq) and reduced representation bisulfite sequencing (RRBS) was performed on brown adipose tissue (BAT) which provide the fundament for the identification of differentially methylated positions (DMP) and regions (DMR) as well as differentially expressed genes (DEG) in three models: *COLD_chow_*, comparing 8° vs 30°C mice on chow diet; *COLD_HFD_*, comparing 8° vs 30°C mice on HFD diet and *ΔCOLD_diet_*, comparing *COLD_chow_* vs *COLD_HFD_*. Differentially methylated and expressed genes (DMEGs; circos plot shows a representative example) were identified for all comparisons based on those DMPs which correlated significantly with differentially expressed genes (DEGs; the heatmap shows a representative example). DMEGs were taken forward for pathways enrichment analyses and to identify differentially regulated epigenetic regulators using the EpiFactors Database. The figure was created with BioRender.

## 1. Introduction

Brown adipose tissue (BAT) is a specialized fat tissue that is rich in mitochondria and promotes non-shivering thermogenesis by expressing the *uncoupling protein 1 (UCP1).* Unlike white adipose tissue, it burns calories to produce heat and therefore plays a key role in energy regulation and controlling metabolic health. Increasing energy expenditure by BAT activation is an intriguing therapeutic approach to combat the overwhelming obesity pandemic, either alone or to complement the current pharmacotherapy that mainly addresses energy intake based on the incretin-mimetic poly-agonist class of drugs [1]. With this in mind it is not surprising that a lot of research was conducted to understand the molecular underpinnings of BAT regulation specifically addressing environmental cues. Cold exposure is the most powerful inducer of BAT activation leading to the upregulation of thermogenic gene program and adrenergic receptor-mediated activation of lipolysis and metabolism. BAT activation also occurs post-prandially, especially after acute overfeeding, to trigger diet-induced thermogenesis [2,3]. However, this compensatory component of energy-expenditure is impaired during chronic overfeeding, a phenomenon that was termed adaptive thermogenesis, and is believed to further drive weight gain and obesity [2,4,5].

In any case the mechanistical switch enabling the high plasticity of BAT changing from passive thermoneutral to active thermogenic states may be regulated by epigenetic mechanisms that are affected by, e.g. temperature and diet [6]. Indeed, studies have shown changes in DNA methylation for key BAT regulatory genes such as *Ucp1, PR domain containing 16* (*Prdm16*) or *PPARG coactivator 1 alpha* (*Ppargc1a*) , altering their expression level and impacting BAT function [7,8]. Moreover, during the development of precursor cells into major brown adipocytes, DNA methylation patterns change and lineage commitment towards a thermogenic adipocyte is epigenetically controlled [8]. The major relevance of the ambient temperature in this context has just recently been implied, as cold exposure of parents at conception is related to increased brown adipogenesis and thermogenesis along with reduced obesity risk in the offspring [9]. Mechanistically, sperm DNA methylation is altered in response to cold exposure which leads to adaptations in BAT activity of the offspring [10,11]. In addition, DNA methylation in key genes of thermogenesis and fatty acid oxidation, such as *acetyl-CoA acyltransferase 2* (*Acaa2*) and *acyl-CoA synthetase long chain family member 1* (*Acsl1*), is increased in BAT of offsprings from mice fed a high fat diet [12]. These studies strongly imply the relevance of cold, diet and obesity on the transgenerational modulation of BAT function. However, these environmental cues, either alone or in combination, also trigger and control BAT plasticity during adulthood. Rats with a blockade in afferent vagal nerve signaling that triggered by high-fat diet showed a reduced BAT activation after cold exposure [13]. The relevance of diet for regulating DNA methylation in later life comes with nutritional compounds such as folate or methionine as critical components of the one-carbon metabolism, which are classically enriched in vegetable-rich diets but diminished in high fat diets.

However, the interaction of especially cold exposure and high fat diet on BAT’s epigenetic machinery is less understood. In this study, we aimed to address DNA methylation alterations in response to temperature and diet, and link this to changes in gene expression of BAT in mice on a genome-wide level. We explored the interactive impact of diet and cold exposure on the epigenetic regulation of BAT gene expression. Using a stringent bioinformatic workflow we focused on differentially methylated and expressed genes (DMEGs) by taking forward only candidates whose differential methylation signal correlates to the gene expression level in order to provide only top functional relevant candidates. Providing a profound overview over the impact of DNA methylation on genes expression affected by cold and diet is of strong relevance to understand adult BAT plasticity and adaptive thermogenesis and provides the fundament for new obesity treatment strategies.

## 2. Material and methods

## 2.1. Animal studies

Male C57BL/6NTac mice (Taconic Bioscience, Lille Skensved, Denmark) were housed in pathogen-free facilities at 23°C on a 12h light/dark cycle at the Sächsische Inkubator für Klinische Translation (SIKT), Leipzig. Starting from five weeks of age, mice were either fed a standard chow diet (EV153, 3.3% from fat, Ssniff, Soest, Germany) or a high fat diet (HFD, E15742, 60 kJ% from fat, Ssniff) with ad libitum access to food and water. At 11 (chow) or 22 (HFD) weeks of age, mice were adapted to single housing in rodent climate chambers (MKKL1200, Flohr Instruments, Netherlands) for 5 days, before randomization into four groups (each N = 8 mice) and housed either at thermoneutrality (30°C) or in the cold (8°C) for 7 days. After a 4-hour fasting period, body weight and rectal temperature were measured. BAT and tail surface temperatures were measured by thermal imaging (VarioCAM®, InfraTec, Dresden, Germany). Mice were sacrificed and tissue samples of intrascapular brown adipose tissue (BAT) were collected, snap frozen in liquid nitrogen and stored at -80°C before further analyses. All animal experiments were approved by the local authorities of the Free State of Saxony, Germany (Landesdirektion Leipzig: TVV51/20), as recommended by the responsible local animal ethics review board.

### 2.2. RNA sequencing of brown adipose tissue

RNA from BAT was isolated using RNeasy Lipid Tissue Mini kit (Qiagen, Hilden, Germany) as specified by the manufacturer. RNA sequencing (RNAseq) was performed by the core unit DNA-Technologien, Medical Faculty Leipzig. 50ng of total RNA were depleted of ribosomal RNA using the NEBNext rRNA Depletion Kit v2 (NEB, USA) according to the manufacturer’s instructions. Depleted RNA was transcribed using SuperScript IV reverse transcriptase (ThermoFisher Scientific, USA) for 2h at 55°C. After second strand synthesis (TargetAmp kit, EPICENTRE, USA), the DNA was fragmented using the Illumina Tagment DNA TDE1 Enzyme and Buffer Kits, which fragments DNA and inserts partial sequencing adapter sequences. Final PCR amplification of the libraries was done using KAPA HiFi HotStart Library Amplification Kit with unique dual indexing by IDT for Illumina Nextera DNA Unique Dual Indexes Sets. The barcoded libraries were purified and quantified using Qubit Fluorometric Quantification (ThermoFisher Scientific, USA). Size distribution of the library DNA was analyzed using the FragmentAnalyzer (Agilent, USA). Sequencing of 2x150 bp was performed with NovaSeq sequencer (Illumina, USA) according to the manufacturer’s instructions, with a read depth of 20 million reads per sample.

### 2.3. Reduced representation bisulfite sequencing in brown adipose tissue

Genomic DNA from BAT samples were isolated using the DNeasy blood and tissue kit according to the manufacturer’s instructions (Qiagen, Hilden, Germany). Reduced representation bisulfite sequencing (RRBS) library preparation was performed on 100 ng DNA using the Zymo-Seq RRBS Library Kit (Zymo Research, Freiburg, Germany) by the core unit DNA-Technologien, Medical Faculty Leipzig. Briefly, DNA was digested with 20U/µl MspI for 4h followed by RRBS adapter ligation using 400U/µl T4 DNA ligase and gap filling with 2U/µl Taq DNA polymerase and 10mM 5-methylcytosine dNTP Mix. Bisulfite conversion was performed with lightning conversion reagent, and the final samples were amplified using 5µM of the unique dual index Primer Set (indexes 1-12) supplied by the manufacturer. Quality control was performed on the Bioanalyzer (Illumina, USA) prior paired-end sequencing on the NovaSeq6000 platform (Illumina, USA) were 20 million reads per sample were obtained.

### 2.4. RNAseq data preprocessing

Raw RNAseq reads where quality-filtered and adapter-trimmed using Trimmomatic (v0.39) [14], discarding reads shorter than 36 bp. Afterwards, reads were aligned to GRCm38.p6 reference assembly using STAR (v2.7.8a) [15], allowing for a maximum of 50 multi-mapped reads. Gene counts were quantified with featureCounts from the subread software package (v2.0.1) [16] with multimappings counted by fractions.

### 2.5. Differential gene expression analysis

Transcripts without gene annotation and genes with a total count sum <10 were removed resulting in 18k genes for subsequent gene expression analysis. Variance stabilization transformation (VST) was applied to the filtered gene count matrix using the R package DESeq2 (v1.42.1) [17]. A principal component analysis (PCA) was conducted to verify clustering of diet (HFD, chow) and temperature (8°C, 30°C) groups within the gene expression signals. Three samples were identified as outliers which did not cluster in their respective group according to the first two principal components and were subsequently removed from the analysis. This resulted in group sizes of 7-8 mice reflecting all diet-temperature combinations (N_total_ = 29; N_HFD_30°C_ = 7; N_HFD_8°C_ = 7; N_chow_30°C_ = 7; N_chow_8°C_ = 8). Afterwards, differential gene expression (DGE) analysis was performed with DESeq2. Three comparisons were conducted: chow-fed mice at 8°C versus those at 30°C (*COLD_chow_*), HFD-fed mice at 8°C versus those at 30°C (*COLD_HFD_*), and an interaction model comparing *COLD_HFD_* versus *COLD_chow_* contrasts (*ΔCOLD_diet_*). DEGs with a false discovery rate (FDR) < 0.05 were considered statistically significant. DEG lists from each contrast where subsequently matched with entries in the EpiFactors Database (v2.0, https://epifactors.autosome.org, accessed on 10.09.2024) to investigate diet-specific methylation regulators.

### 2.6. RBBS data processing

Raw RRBS reads from N = 29 mice (N_HFD_30°C_ = 7; N_HFD_8°C_ = 7; N_chow_30°C_ = 7; N_chow_8°C_ = 8) were processed with the nextflow (v 23.04.3) [18] pipeline nf-core/methylseq (v2.4.0) [19] using the parameters *--genome GRCm38 --non_directional --rrbs --skip_deduplication* and *-- cytosine_report*. In brief, quality filtering and adapter trimming of raw RRBS fastq files was conducted with trimgalore (v0.6.7) [20] before running sequence alignments with Bismark (v0.24.0) [21]. Afterwards, methylation signals are called with the Bismark methylation extractor script including the build-in *coverage2cytosine* module for a genome-wide cytosine methylation report which served as input for downstream analysis. Sex chromosomes were excluded.

### 2.7. Differential methylation analysis

To identify differentially methylated positions (DMPs) and differentially methylated regions (DMRs), differential methylation analysis was conducted for *COLD_chow_*, *COLD_HFD_*, and *ΔCOLD_diet_*, utilizing beta values and the R packages limma (v3.54.0) [22] and RnBeads (v2.16.0) [23] with parameters filtering.low.coverage.masking = TRUE, filtering.high.coverage.outliers = TRUE and filtering.snp = “any”. Based on their chromosomal position and respective annotation, individual methylations can be contextualized within gene bodies or promoter regions (1500 bases upstream and 500 bases downstream of the transcription start site), both of which were considered in our analysis.

### 2.8. Integration of methylation and transcriptome data

After differential analysis of individual omics data, both gene expression and methylation signals were combined to find biological associative effects between transcriptomics and epigenomics. The R package biomaRt (v2.58.2) [24] was used to gather gene loci information for differentially expressed genes (DEGs) and DMRs in gene and promotor context. Beta methylation values were transformed to M-values using RnBeads. For each DEG, the corresponding DMR was selected based on the alignment of gene IDs. M-values of all significantly DMPs overlapping with the DMR were considered using the bedR (v1.0.7) [25] and the GenomicRanges (v1.54.1) [26] R packages. Pearson correlation was then applied to find associations between DEG expression values and M-values of each intersecting DMP using the multtest.cor function from RVAideMemoire (v0.9-83-7) [27]. All DMPs with significant absolute Pearson correlations >0.5 between expression and methylation signal (p <0.05 without adjustment for multiple testing) were defined as differentially methylated and expressed genes (DMEGs). If a DMEG is associated to both promoter and gene body region, it was assigned to the DMR with the highest significance. This step was parallelized with the foreach package (v1.5.2) [28]. The final list of DMEGs is comprised of genes with significantly regulated expression signal (absolute log_2_ fold change (FC) >0.5) and methylation signal (absolute log_2_ FC >1). Since several DMRs or DMPs can be differentially methylated within a gene, several DMEGs were generated for such genes.

Subsequently, functional analysis of DMEGs was done with the use of QIAGEN Ingenuity Pathway Analysis software (QIAGEN Inc., https://digitalinsights.qiagen.com/IPA, Winter Release Q4 2024) [29] covering the catalog for disease and bio functions and canonical pathways. The database query was limited within the IPA software to relationships with adipose tissue or adipocytes in mice that are experimentally observed. Enrichments are reported with p after Bonferroni-Holm (BH) adjustment and activation z-scores. Enrichments with BH corrected p <0.05 are considered significant.

### 2.9. In vitro validation experiments

#### 2.9.1. Immortalized and primary brown adipocytes

Immortalized (imBA [30]) and primary brown adipocytes were cultured and differentiated as previously described [31]. At day 10, cells were harvested and subjected to further analyses.

### 2.9.2. In vitro manipulation of cellular DNA methylation levels

To analyze the effect of DNA methylation status on gene expression in imBA and primary brown adipocytes, pre-adipocytes were treated with indicated concentrations of 5’aza-2’-deoxycytidine (Aza) for 48h and subsequently differentiated [31]. Differentiated adipocytes at day 6 were treated with indicated concentrations of S-adenosylmethionine (SAM, New England Biolabs) for 48h. Differentiated adipocytes were harvested and subjected to further analysis.

### 2.9.3. Quantitative real-time-PCR (qPCR)

RNA from imBA and human primary SAT cells was isolated using RNeasy Lipid Tissue Mini kit. qPCR was performed using the LightCycler System LC480 and LightCycler-DNA Master SYBR Green I Kit (Roche, Mannheim, Germany). Adipocyte gene expression of candidate genes was calculated by ΔΔCT method and normalized to *Nono* or *36b4* levels as indicated. Primer sequences are listed in (Suppl. Table 1).

### 2.10. Statistical analysis

Analyses were performed under R version 4.3.2, except for mouse phenotyping data that was analyzed using GraphPad Prism 10 (GraphPad, San Diego, CA, USA). Statistical tests are stated in the figure legends. If not stated otherwise, adj. p <0.05 were considered statistically significant.

## 3. Results

### 3.1. Phenotyping of chow and high fat diet-fed mice held at thermoneutrality or exposed to cold

Groups of 8 mice per diet were held at thermoneutrality (30°C) or in the cold (8°C) for 7 days (Figure 1A). Before the temperature challenge, mice were held at 23°C with HFD-fed mice presenting significantly higher body weights than chow fed mice (48.2 g vs 28.3 g, Figure 1B). In the cold, mice on chow diet maintained a steady body weight, while mice on HFD lost about 10% body weight (Figure 1B, C). All mice in the cold showed a significant drop in body temperature during the first days, but this loss in temperature was significantly higher in mice on HFD (−0.5°C vs -2°C after d1, Figure 1D). After 7 days, mice on chow diet recovered their body temperature to normal levels, while mice on HFD still had significantly lower temperatures (−1°C) (Figure 1D, E). Under thermoneutrality, mice on HFD had significantly higher BAT surface temperatures, consistent with increased diet-induced thermogenesis, but BAT surface temperatures were not different after cold exposure (Figure 1F, G). Extended phenotyping data is presented in Suppl. Figure 1.

**Figure 1:**
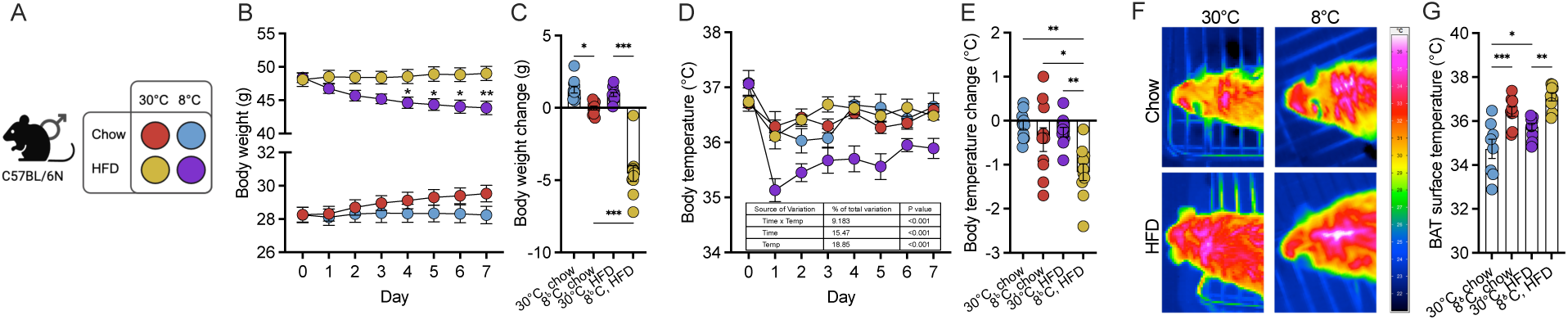
Phenotyping of chow and HFD-fed mice housed at 30°C or 8°C. (A) Schematic overview and corresponding color code of the cohort mouse groups stratified by dietary (chow, HFD) and environmental (thermoneutrality vs cold) conditions. (B) Body weight development and (C) body weight change in chow and HFD-fed mice housed at thermoneutrality (30°C) or in the cold (8°C) for 7 days. Body temperature (D) and temperature change (E) in chow and HFD-fed mice housed at thermoneutrality (30°C) or in the cold (8°C) for 7 days. (F) Thermal images from brown adipose tissue (BAT) and (G) BAT surface temperature in chow and HFD-fed mice housed at thermoneutrality (30°C) or in the cold (8°C) for 7 days. N = 7-8 per condition. (B, D) Statistical significance was evaluated by two-way ANOVA with uncorrected Fischer’s LSD, (E) Šídák’s post-hoc test, (C, G) or one-way ANOVA with Tukey’s post-hoc test. p <0.05 (*), p <0.01 (**), p <0.001 (***). Scale bar: 100 µm.

### 3.2. Obesity impairs core transcriptome remodeling of BAT in response to cold

Gene expression data was used from all mice samples that also had methylation signal data available. After removal of outliers, N = 29 samples (N_HFD_30°C_ = 7; N_HFD_8°C_ = 7; N_chow_30°C_ = 7; N_chow_8°C_ = 8) remained for subsequent differential gene expression and methylation analysis. Analysis of VST normalized RNAseq expression counts shows clear clustering of all diet (chow, HFD)- and temperature (8°C, 30°C)-specific subgroups in the principal component analysis (Figure 2A), providing a solid basis for subsequent differential gene expression analysis. Moreover, temperature groups are separated by principal component 1 describing 33% of variance while diet groups are rather distinguished by principal component 2 covering 16,8% of total variance demonstrating an overall stronger impact of temperature compared to diet on overall gene expression patterns. Mice on HFD are older compared to chow fed mice. The age gap of 11 weeks could potentially interfere and explain a variance in the principal component analyses which was not addressed here.

**Figure 2:**
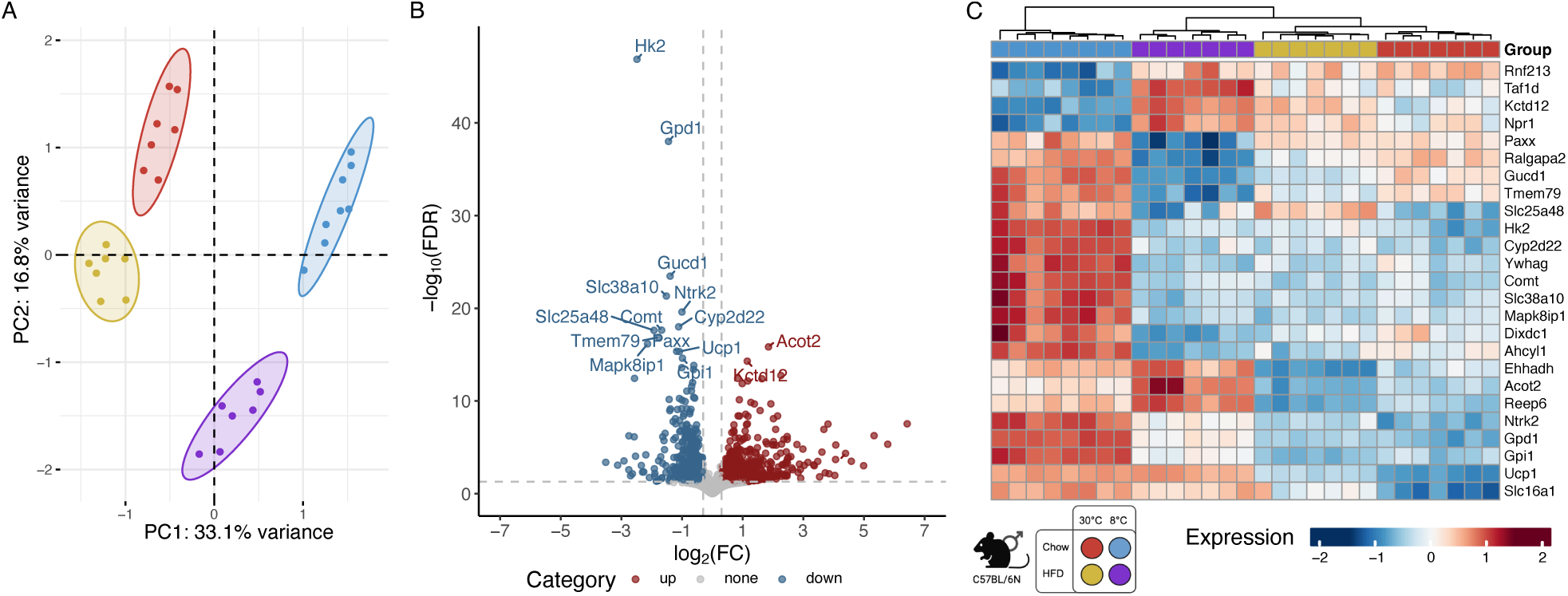
RNA sequencing-based gene expression data revealse distinct clusters of treatment groups. (A) Principal component analysis separates temperature and diet. All considered subgroups reveal distinct expression profiles and cluster according to the first two principal components. (B) Volcano plot highlights most significantly (false discovery rate (FDR) <0.05) regulated genes under *ΔCOLD_diet_* condition. (C) Heatmap of expression values shows that the subgroups can be clearly separated by the top significant regulated genes. Abbreviations: HFD: high fat diet; PC: principal component.

To dissect BAT’s physiological response to cold normal-weight control mice, we compared chow-fed mice held in the cold (8°C) with mice held at thermoneutrality (30°C) for one week. We will refer to this condition pair as *COLD_chow_*. Additionally, to assess the effects of obesity on cold-induced BAT activation, we compared HFD-induced obese mice held in the cold (8°C) with mice held at thermoneutrality (30°C) for one week. This comparison will be referred to as *COLD_HFD_*. Differential gene expression analysis (FDR <0.05) resulted in 7,089 DEGs for the *COLD_chow_* (Suppl. Figure 2A, B; Suppl. Table 2) and 5,249 DEGs for the *COLD_HFD_* (Suppl. Figure 2C, D; Suppl. Table 3) condition pairs. Reflecting the dominating effect of temperature on BAT activity, the majority of the significant DEGs overlap between the *COLD_chow_* and *COLD_HFD_* conditions are regulated similarly (N = 3,786; *COLD_chow_*: 53%; *COLD_HFD_*: 72%). Furthermore, 45% and 26% are specifically regulated in *COLD_chow_* and *COLD_HFD_*, respectively, while a total of 92 DEGs are contrarily regulated (Suppl. Figure 4A).

To benchmark our *COLD_chow_* results with previously published data, we overlapped our DEGs with those reported by Taylor et al. (2024) [32], who performed a comparable study in chow-fed male mice exposed to severe cold (8 °C) versus thermoneutrality (28 °C) (Suppl. Figure 3; Suppl. Table 2). We found that 80% of our upregulated and 52% of our downregulated genes overlapped with their DEGs, demonstrating a strong concordance across both directions in these independent analyses.

Additionally, to investigate how diet affects BAT activation in response to cold exposure, we employed an interaction model (Δ*COLD_diet_*) that compares the difference between both *COLD_HFD_* and *COLD_chow_* contrasts, rather than relying solely on simple group comparisons. This approach enabled us to assess the combined and potentially synergistic or antagonistic effects of diet and environment on BAT function [33]. Although an age gap of 11 weeks is indicated, mice on chow or HFD are in a comparable age-range regarding BAT function and diet-independent body weight development [34–37]. In total, we observed 1,364 genes (FDR <0.05) which are differentially expressed between *COLD_HFD_* and *COLD_chow_* (Figure 2B, C; Suppl. Table 4). Since the most significant DEGs are downregulated in the *COLD_HFD_* condition compared to *COLD_chow_*, this highlights a potential mechanism by which obesity may impair thermogenic capacity and energy expenditure, suggesting that a high-fat diet disrupts the normal activation of BAT in response to cold exposure.

### 3.3. Associations of gene expression and methylation signals underscore the influence of diet on gene regulatory mechanisms

To explore the associative effects between transcriptomics and epigenomics, we correlated all expression signals of DEGs with significant DMPs found within the DMRs of the corresponding genes. This approach allowed us to define a set of differentially methylated and expressed genes (DMEGs), using specific criteria: a Pearson correlation coefficient greater than ± 0.5, DEGs with a log_2_ FC greater than ± 0.5 (adj. P <0.05), and DMPs with a log_2_ FC greater than ± 1 (p <0.05). Importantly, we set a higher threshold for DMPs and focused on DMEGs located in both the promoter and gene body regions, ensuring that our analysis captures the most relevant regulatory elements influencing gene expression. To verify our findings, we analyzed the effect of DNA-demethylation (using Aza) or hypermethylation (using SAM) on the expression of selected genes in mouse adipocytes *in vitro* (Suppl. Figure 5). The observed effects of DNA methylation and the manipulation on gene expression were in line with the results of our analysis correlating gene expression and methylation and confirmed epigenetic control of gene expression in adipocytes.

In the *COLD_chow_* control comparison, we identified a total of 3,142 DMEGs among 21% (N = 1,524) of all DEGs. The data showed a balanced distribution of DMEGs, with 55% located within gene bodies and 45% found in promoter regions (Figure 3A; Suppl. Table 5), indicating that methylation can affect gene regulation through various mechanisms, such as promoter accessibility or splice site manipulation. As illustrated in Figure 3B, DMEGs within the top 100 DMRs (ranked by absolute log_2_ FC) are predominantly found in promoter regions and show distinctly higher prevalence on chromosome 11.

**Figure 3:**
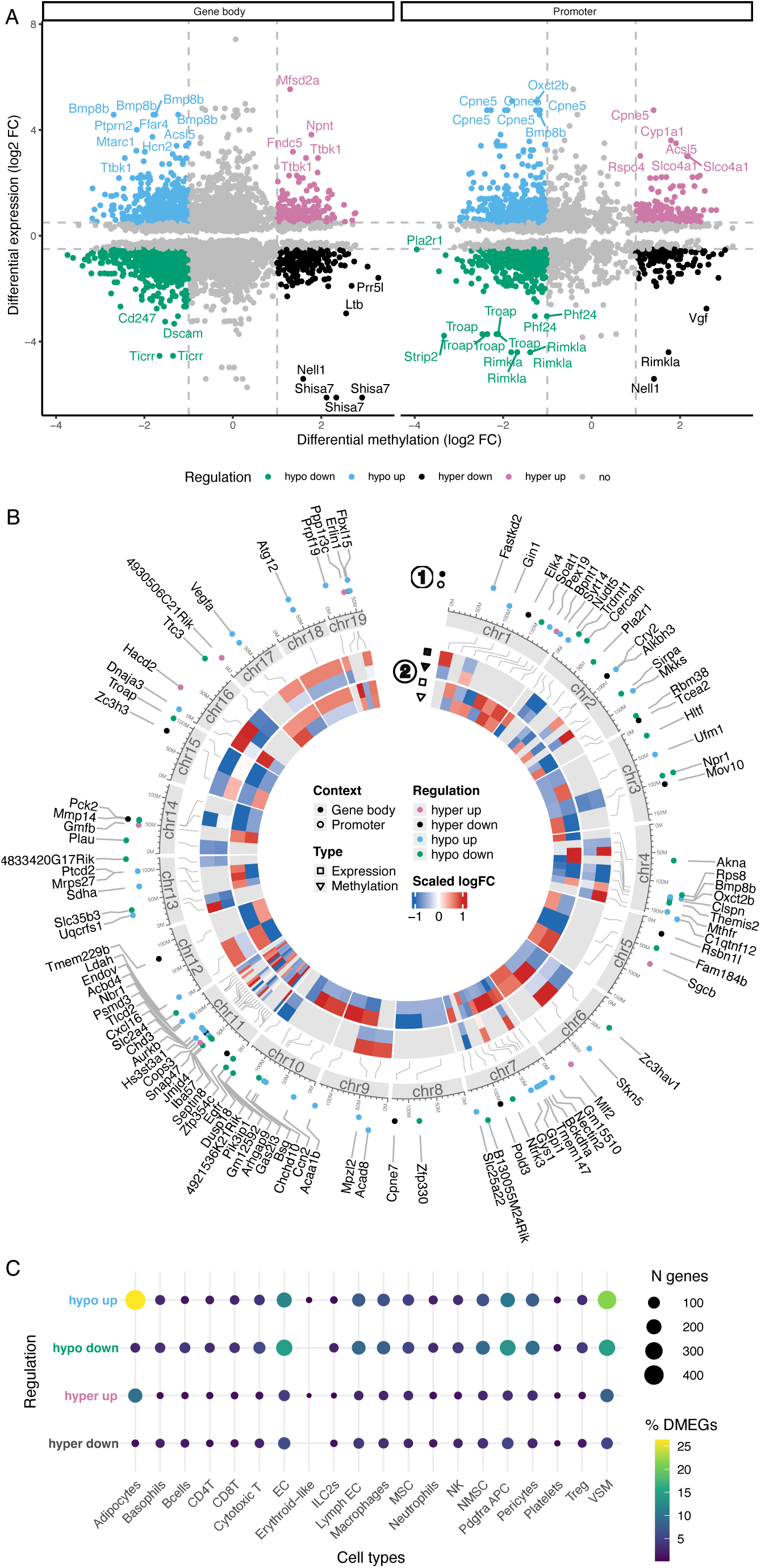
Multi-layered insights into DMEGs under *COLD_chow_* condition. (A) The scatter plot displays the log_2_ fold changes (FC) in gene expression and DNA methylation signals for all differentially methylated and expressed genes (DMEGs) in both gene body and promoter contexts under the *COLD_chow_* condition. Colors indicate the direction of regulation. DMEGs with absolute log_2_ FC >0.5 in expression and >1 in methylation signals are highlighted as significant, while DMEGs in the 5% and 95% quantile of both differentially methylated position (DMP) and differentially expressed gene (DEG) log_2_ FC signals are labeled. (B) The circos visualization illustrates the DMEGs covering the top 100 significant differentially methylated regions (DMRs; ranked by log_2_ FC). This plot integrates multiple layers of information from outer to inner ring: Dot colors indicate the direction of regulation, while dot positions denote whether the DMR of the DMEG is located in the gene body or promoter context (1); a stylized ideogram of chromosomes provides a genomic reference frame; and a heatmap displays scaled log2 FC for both gene expression and DNA methylation signals in gene body and promoter contexts for each DMEG (2). (C) Overview of assigned cell types to the DMEGs, categorized as hypo up, hypo down, hyper up, and hyper down. The color scale reflects the percentage of DMEGs, and the size of the dots reflects the absolute number of DMEGs.

To investigate which cell types might drive DNA methylation changes in BAT, we overlapped DMEGs from the *COLD_chow_* condition with cell-type-specific marker genes derived from single-cell RNAseq (scRNAseq) of BAT from male mice exposed to cold (5 °C) versus thermoneutrality (30 °C) [38]. This analysis revealed that DMEGs were distributed across multiple cell types, with the highest proportions in vascular smooth muscle cells (VSM, 43%), adipocytes (35%), endothelial cells (EC, 31.8%), and stromal populations, including *Pdgfra*⁺ adipocyte progenitor cells (APCs, 28.1%) and pericytes (20.9%). In contrast, immune cells contributed more modestly, with macrophages at 18.2%, NK cells at 9.3%, and T cell subsets (CD4⁺, CD8⁺, Treg) ranging from 5–9%. Across nearly all cell types, most DMEGs belonged to the hypo up category - genes exhibiting hypomethylation accompanied by increased expression. This pattern was particularly pronounced in adipocytes (26.4%) and vascular populations, including VSM (21.6%) and EC (11.9%). Hyper up DMEGs were most notable in adipocytes (10%), whereas hypo down DMEGs were enriched in vascular and stromal populations, including VSM (15%), EC (14.9%), *Pdgfra⁺*APCs (13.6%), and to a lesser extent in adipocytes (2.8%). Conversely, the fractions of hyper down DMEGs were relatively small, ranging from 1-6% depending on the cell type (Figure 3C; Suppl. Table 5).

Additionally, we detected 1,886 DMEGs from 19% (N = 1001) of all DGEs in the *COLD_HFD_* condition. Similar to the *COLD_chow_* condition, 60% of the DMEGs are located within gene bodies and 40% in promoter regions (Figure 4A; Suppl. Table 6); however, the most highly regulated DMEGs do not exhibit significant enrichment in the promoter regions (Figure 4B).

**Figure 4:**
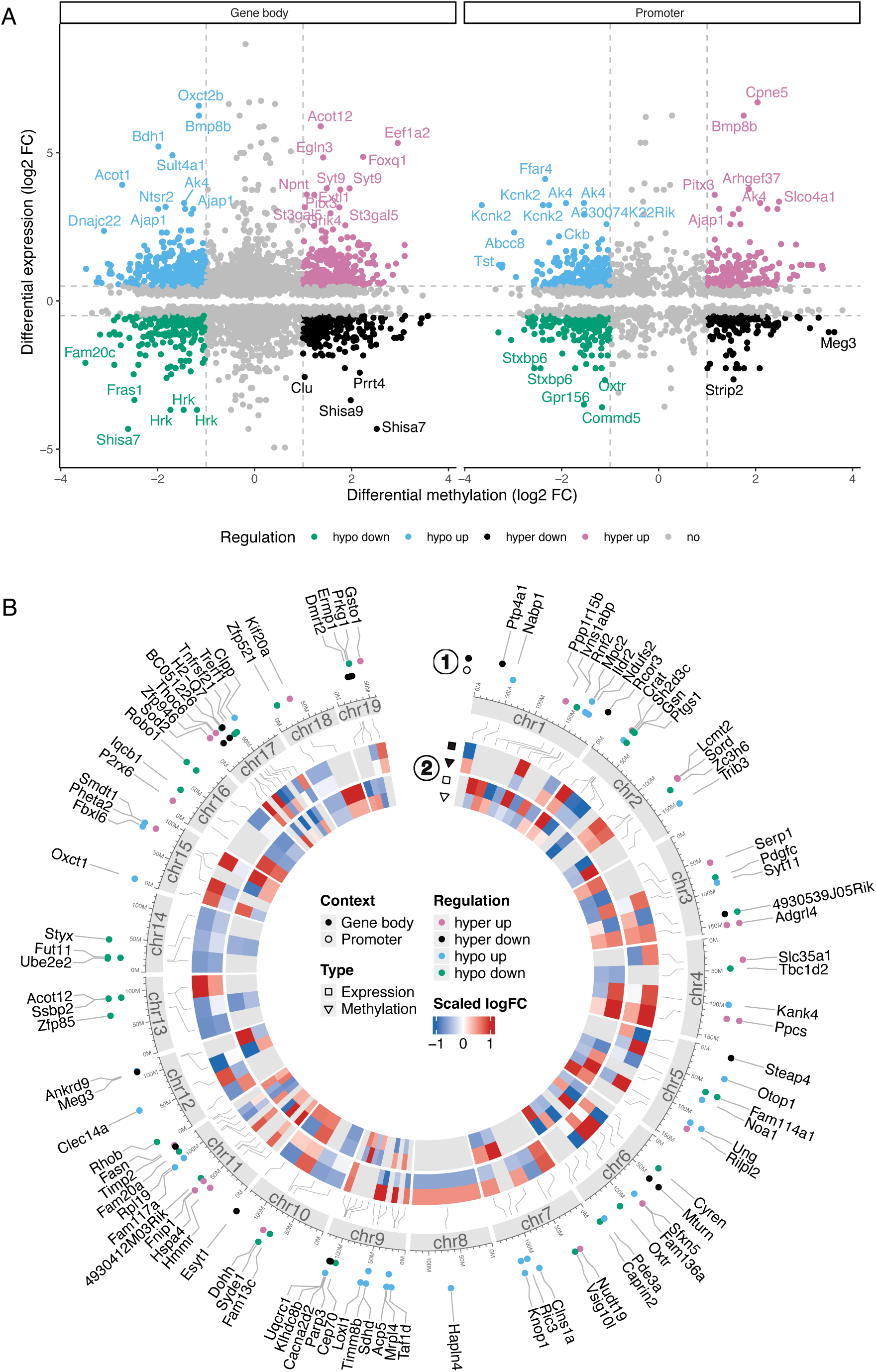
Multi-layered insights into DMEGs under *COLD_HFD_* condition. (A) The scatter plot displays the log_2_ FC in gene expression and DNA methylation signals for all DMEGs in both gene body and promoter contexts under the *COLD_HFD_* condition. Colors indicate the direction of regulation. DMEGs with absolute log_2_ FC >0.5 in expression and >1 in methylation signal are highlighted as significant, while DMEGs in the 5% and 95% quantile of both DMP and DEG log_2_ FC signals are labeled. (B) The circos visualization illustrates the DMEGs covering the top 100 most significant DMRs (ranked by log_2_ FC). This plot integrates multiple layers of information from outer to inner ring: Dot colors indicate the direction of regulation, while dot positions denote whether the DMR of the DMEG is located in the gene body or promoter context (1); a stylized ideogram of chromosomes provides a genomic reference frame; and a heatmap displays scaled log_2_ FC for both gene expression and DNA methylation signals in gene body and promoter contexts for each DMEG (2).

Although we previously observed that the majority of significant DEGs overlapping between the *COLD_chow_* and *COLD_HFD_* conditions are regulated in a similar direction (Suppl. Figure 5), we identified a contrasting pattern among the DMEGs. The gene expression of DMEGs shows consistent regulatory patterns, with only 245 (*COLD_chow_*: 16%; *COLD_HFD_*: 24%) genes being upregulated and 159 (*COLD_chow_*: 10%; *COLD_HFD_*: 16%) genes being downregulated in both conditions (Suppl. Figure 4B; Suppl. Figure 6). In contrast, 73% and 60% are specifically regulated in the *COLD_chow_* and *COLD_HFD_* conditions, respectively, while only one gene (*guanylyl cyclase domain containing 1* (*Gucd1*)) is regulated in the opposite direction (Suppl. Figure 4B; Suppl. Figure 7). This suggests that the response to temperature conditions is strongly dependent on the specific environment and that methylation changes may be purposefully employed to adapt to different metabolic or physiological demands.

Our previous findings revealed pronounced differences in the regulatory patterns of DMEGs between the *COLD_chow_* and *COLD_HFD_* conditions by emphasizing the direction of regulation. To further elucidate the nuances of diet-specific regulation, we utilized the Δ*COLD_diet_* contrast to detect more subtle differences in regulatory effects including regulation strength, directly identifying diet-related differences in DMEGs associated with BAT activation induced by cold exposure (Figure 5; Suppl. Table 7). For the Δ*COLD_diet_* contrast, we detected 65 DMEGs from only 4% (N = 56) of all DEGs. Interestingly, these are primarily enriched in the gene bodies (69%), suggesting that the regulation of these genes may be more influenced by post-transcriptional mechanisms in response to cold exposure.

**Figure 5:**
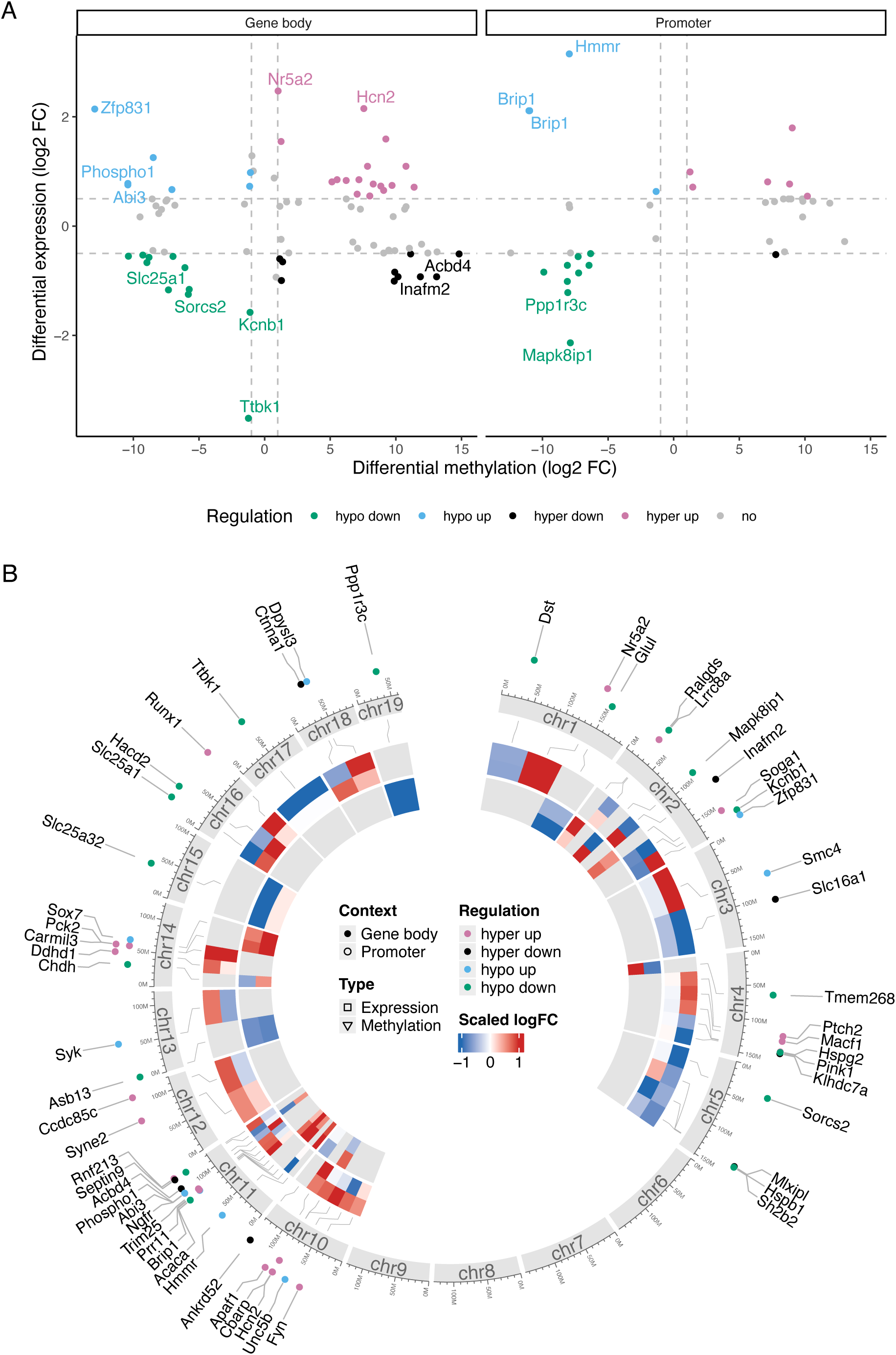
Multi-layered insights into DMEGs under *ΔCOLD_diet_* condition. (A) The scatter plot displays the log_2_ FC in gene expression and DNA methylation signals for all DMEGs in both gene body and promoter contexts under the *ΔCOLD_diet_* condition. Colors indicate the direction of regulation. DMEGs with absolute log_2_ FC >0.5 in expression and >1 in methylation signals are highlighted as significant, while DMEGs in the 5% and 95% quantile of both DMP and DEG log_2_ FC signals are labeled. (B) The circos visualization illustrates all detected DMEGs in their respective DMRs. This plot integrates multiple layers of information from outer to inner ring: Dot colors indicate the direction of regulation, while dot positions denote whether the DMR of the DMEG is located in the gene body or promoter context (1); a stylized ideogram of chromosomes provides a genomic reference frame; and a heatmap displays scaled log_2_ FC for both gene expression and DNA methylation signals in gene body and promoter contexts for each DMEG (2).

### 3.4. Adaptive divergence in thermogenic signaling reflects metabolic stress in obese BAT

For further characterization of the reported DMEGs, pathway enrichment analysis was performed using the IPA software. The DMEGs were categorized into four groups: those exhibiting similar regulation in both *COLD_chow_* and *COLD_HFD_* condition pairs, and those that are uniquely regulated in each individual condition (Δ*COLD_diet,_ COLD_chow_*, and *COLD_HFD_*). Querying the catalog for significant (BH-adjusted p <0.05) canonical pathways (Figure 6; Suppl. Table 8) shows similar activation of pathways with respect to lipid metabolism except for ketolysis which is only significantly activated in the *COLD_HFD_* condition. Cell signaling and energy expenditure seem to be activated or inhibited more differentially between *COLD_chow_* and *COLD_HFD_* condition pairs. For example, the process of white adipose tissue browning is significantly inhibited in *COLD_HFD_* but shows (unsignificant) activation in *COLD_chow_*. Additionally, type II diabetes mellitus signaling is significantly activated under *COLD_HFD_* considerations but tends to be (insignificantly) inhibited under *COLD_chow_*.

**Figure 6:**
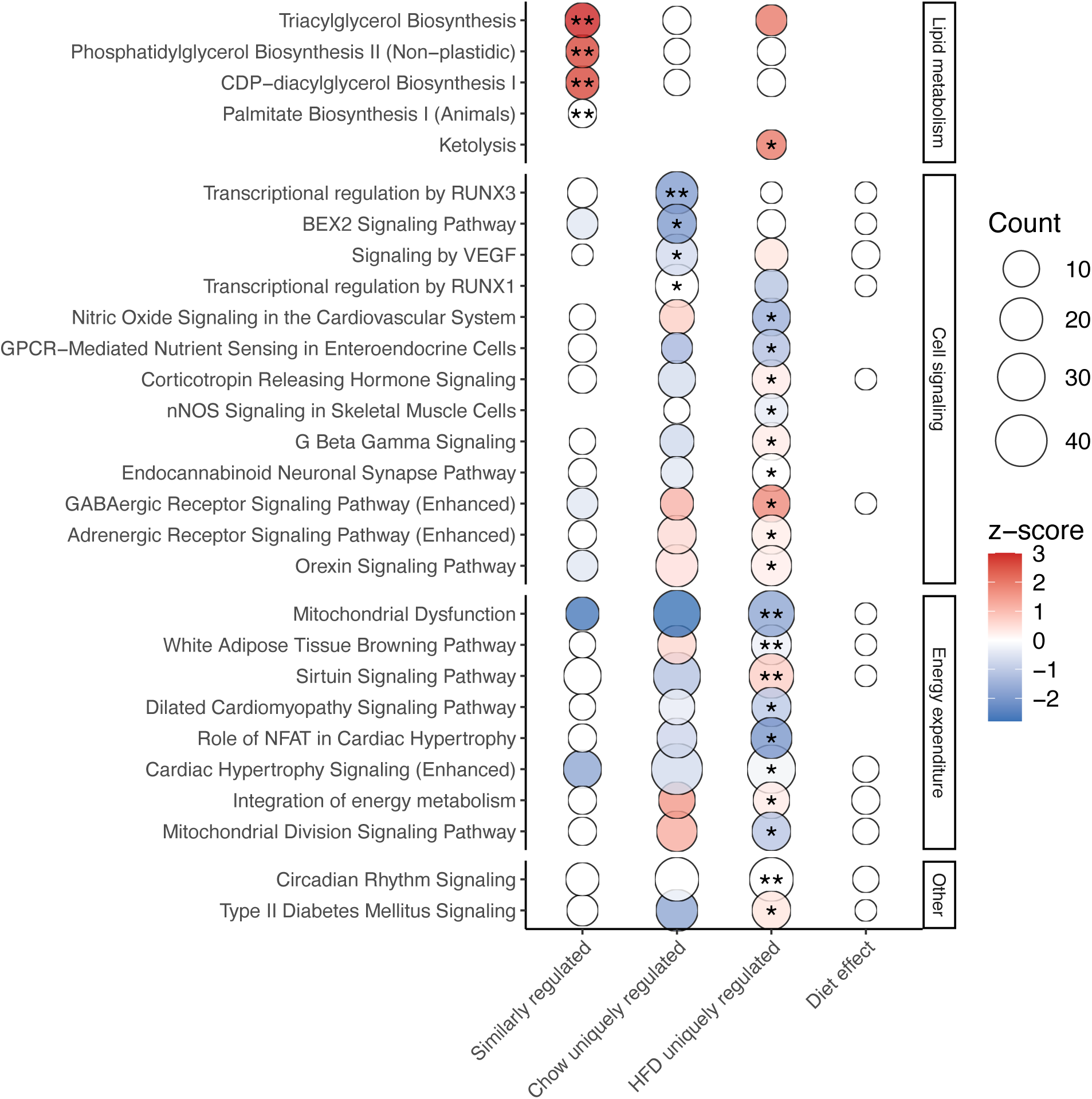
Pathway enrichment analysis of DMEGs for canonical pathways, diseases, and biological functions. Ingenuity Pathway Analysis (IPA) with enrichments of DMEGs were categorized into four groups: those exhibiting similar regulation in both *COLD_chow_* and *COLD_HFD_* condition pairs, and those that are uniquely regulated in each individual condition (Δ*COLD_diet,_ COLD_chow_*, and *COLD_HFD_*). Top ten statistically significant enrichments are shown for canonical pathways. More then ten enrichments are listed in case of p ties. Circle sizes reflect the number of genes in each enrichment while the color highlights the activation z-score. Significance levels for p after adjustment for multiple testing with Bonferroni-Holm are indicated as p <0.05 (*), p <0.01 (**).

### 3.5. Diet and cold exposure shape epigenetic regulation in BAT

To gain deeper insights into the epigenetic mechanisms that regulate BAT activation and thermogenesis under different dietary conditions, we matched DEGs from each condition pair with the EpiFactors database (v2.0, https://epifactors.autosome.org, accessed on 10.09.2024), with the analysis being restricted to DNA-methylation-related factors, while histone-modifying enzymes, chromatin remodelers and RNA-based epigenetic regulators were omitted. As summarized in Figure 7, we identified eleven differentially expressed methylation regulators for the *COLD_chow_*, seven for *COLD_HFD_*, and four for Δ*COLD_diet_* condition pairs. Notably, five regulators were found to be common to both the *COLD_chow_* and *COLD_HFD_* conditions including *Tet2* (*ten-eleven translocation methylase 2*), *Dnmt3* (*DNA methyltransferase 3*), *Cenpc1* (*centromere protein C*), *Apobec1* (*apolipoprotein B mRNA editing enzyme catalytic subunit 1*) which are all downregulated in *COLD_chow_ and COLD_HFD_*. This overlap indicates a conserved epigenetic mechanism that is essential for the physiological response to cold exposure, regardless of diet. However, the majority of methylation enzymes exhibited unique regulation within each condition pair, highlighting the nuanced regulation of epigenetic mechanisms based on dietary composition. For instance, *Tet3* and *Dnmt1* were differently regulated within the Δ*COLD_diet_* and *COLD_chow_* conditions, while *Mettl4* (*methyltransferase-like protein 4*) and *Apobec3b* showed differential regulation within the Δ*COLD_diet_* contrast.

**Figure 7:**
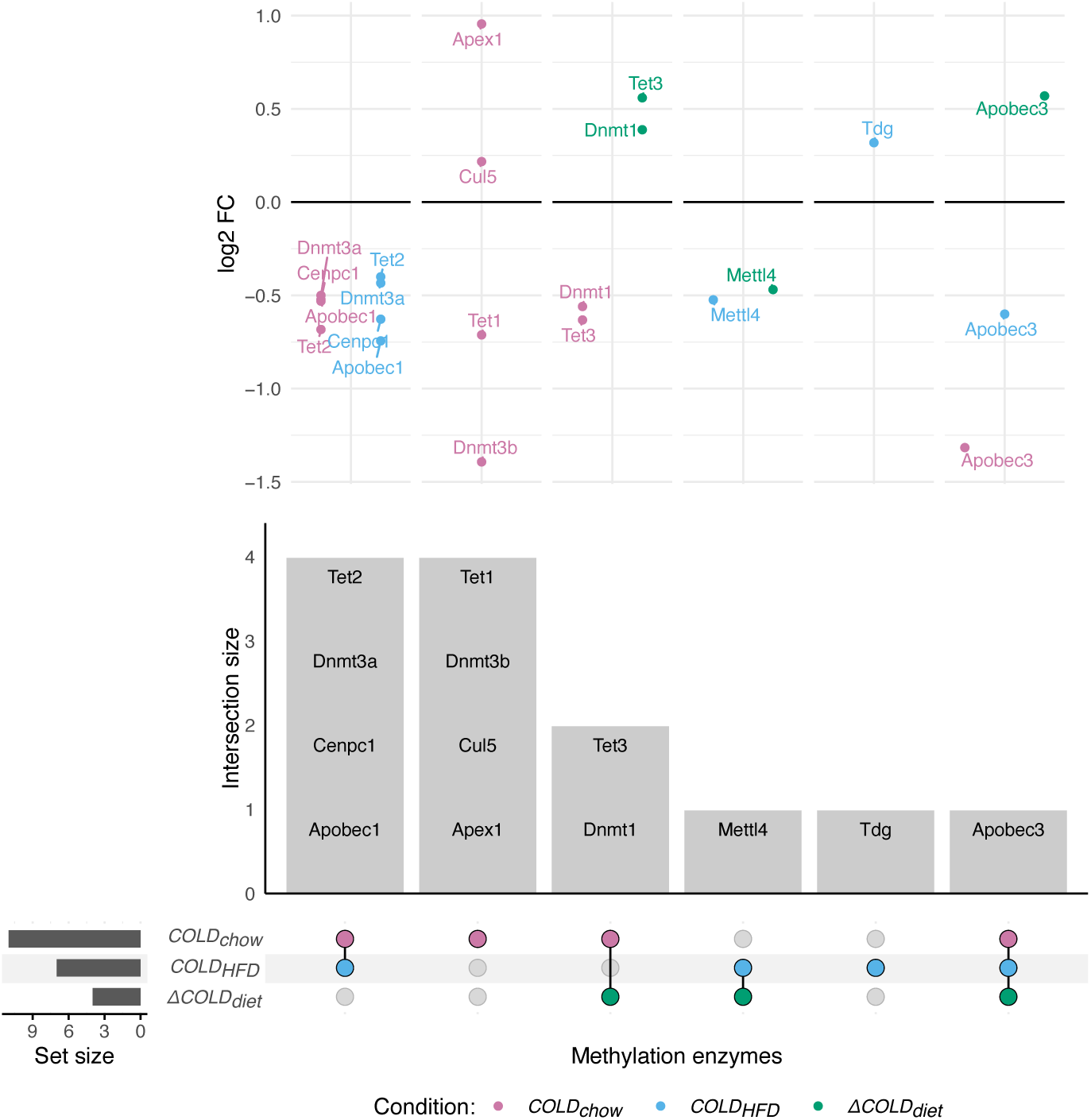
Overlap of methylation regulators in cold-induced BAT activation across dietary conditions. Upset plot displays the overlap of differentially expressed methylation regulators detected for the *COLD_chow_*, *COLD_HFD_*, and Δ*COLD_diet_* condition pairs. The scatterplot shows log2 FC values for each differentially expressed methylation regulator of the respective intersect in each condition highlighted by coloring.

## 4. Discussion

We aimed to explore how environmental factors and DNA methylation may influence BAT function. To investigate potential links between DNA methylation and gene expression, we integrated RNA-seq data with RRBS methylation profiles to identify genes with coordinated changes in expression and methylation (DMEGs) under different dietary and thermal conditions. This approach provides insights into BAT plasticity and adaptive thermogenesis, contributing to a foundational understanding that could inform future obesity-related research and interventions.

Consistent with BAT’s physiological adaptation to cold, our data show that cold exposure robustly increases BAT activity regardless of diet, evidenced by strong transcriptomic responses with 7,089 DEGs in chow-fed and 5,249 DEGs in HFD-fed mice. A substantial overlap (∼3,780 genes) between both dietary groups suggests a conserved “core” thermogenic program that remains inducible under obesogenic conditions. Notably, *Ucp1*, a central mediator of non-shivering thermogenesis, and *Letmd1* (*leucine zipper-EF-hand-containing transmembrane protein domain containing 1*), which is essential for mitochondrial integrity and cold response, were among the top upregulated genes across both diets. Their consistent induction under cold exposure, aligns with previous reports of diet-insensitive thermogenic regulators [39,40]. Beside this shared response, ∼1,360 genes were differentially expressed between the cold-exposed groups (*ΔCOLD_diet_*), with most downregulated in diet-induced obese mice, consistent with a high-fat diet impairing thermoregulatory capacity driven by mitochondrial dysfunction, inflammation, and insulin resistance [41,42]. This functional attenuation under high-fat diet conditions is also reflected among the most strongly regulated DGEs. Downregulation of *Hk2* (*hexokinase 2*), a key glycolytic enzyme, has been linked to reduced glucose uptake [43]. Similarly, reduced *Gpd1* (*glycerol-3-phosphate dehydrogenase 1*) expression, part of the glycerol-3-phosphate shuttle, may reflect compromised lipid metabolism. *Ucp1* stayed cold-inducible in all diets but was lower in HFD-fed mice, showing that thermogenic response is maintained but reduced under metabolic stress. This aligns with reports of reduced BAT activity in obesity despite retained gene expression [39]. Our results of *COLD_chow_* are in line with those of Taylor et al. (2024) [32], who performed a comparable study in chow-fed mice exposed to severe cold (8°C) versus thermoneutrality (28°C). The strong overlap of DEGs highlights the reproducibility of cold-induced BAT transcriptional responses across independent studies and supports the generalizability of our findings.

To identify epigenetically regulated genes, we correlated methylation changes (M-values of DMPs overlapping DMRs) with gene expression levels to define DMEGs. While these correlations suggest potential regulatory links, they do not establish causality, as gene expression is also influenced by other epigenetic modifications and transcription factors [32,44,45]. More comprehensive approaches integrating histone modifications, proteomics, or single-cell epigenomics could further elucidate the complex regulation within heterogeneous BAT cell populations [46]. Compared to generalized linear models with distributional assumptions [47,48], our site-specific correlation approach offers higher sensitivity for detecting local methylation-expression relationships, though it may miss broader regulatory patterns and is more sensitive to noise. Our thresholds for defining significant DMEGs were set at |log FC| >0.5 for expression changes and |log FC| >1 for methylation differences; correlations were considered significant if |r| >0.5 and p <0.05. These cutoffs balance sensitivity and specificity but also limit detection power given our sample size. When DMEGs overlapped promoter and gene body regions, assignment was based on the strongest correlation, which enhances interpretability but may introduce bias due to regional variability or technical factors.

By analyzing DMEGs, ∼3,100 DMEGs (21% of DGEs) were identified in *COLD_chow_*, with a roughly equal distribution between promoter and gene body methylation. In contrast, HFD-fed mice exhibited only ∼1,900 DMEGs (19% of DEGs), with a higher proportion of gene body methylation. As gene body methylation is often linked to transcript fine-tuning rather than activation, this may reflect constrained thermogenic adaptability under obesogenic conditions [49]. Notably, DMEGs were largely condition-specific despite substantial overlap in DEGs, highlighting that diet and cold exposure interact to shape BAT-specific regulatory states, rather than acting independently. These condition-specific methylation-expression relationships suggest that DNA methylation may play an active role in modulating transcriptional plasticity and energy metabolism, enabling BAT to adapt epigenetically to distinct environmental challenges.

To gain further insight into the cellular context of these DNA methylation changes, we mapped DMEGs of the *COLD_chow_* condition onto cell-type-specific markers derived from male mice scRNAseq data of cold-exposed BAT [38]. This analysis revealed that the majority of DMEGs belonged to the hypo up category, particularly in adipocytes, vascular cells, and *Pdgfra⁺* adipocyte progenitors, suggesting coordinated activation of thermogenic programs and vascular remodeling. Hypo down DMEGs were enriched in vascular and stromal populations, indicating selective repression of pathways supporting tissue structure and function. Hyper up DMEGs were primarily observed in adipocytes, whereas hyper down DMEGs were rare (1–6%), implying that DNA methylation mediated silencing plays a limited role in the cold response. The observed cell type-specific DMEG distributions are consistent with prior evidence of increased intercellular signaling among adipocytes, vascular, stromal, and immune cells under cold conditions in chow-fed mice, suggesting an epigenetic basis for the transcriptional remodeling in BAT [38].

Pathway enrichment analysis further supports this adaptive divergence of DMEGs and highlighting the diet-dependent rewiring of thermogenic signaling. Cold exposure activated distinct lipid metabolic pathways (triacylglycerol, phosphatidylglycerol, and CDP−diacylglycerol biosynthesis) in both dietary groups, however, ketolysis was selectively enriched in HFD-fed mice. In line with previous findings that HFD suppresses ketone body utilization for lipogenesis in BAT via downregulation of *acetoacetyl-CoA synthetase* (*Aacs*) [50], our DGE analysis show that *Aacs* was significantly upregulated by cold in both dietary groups, but this response was markedly stronger in chow-fed mice (log_2_ FC: 1.77) than in HFD-fed mice (log_2_ FC: 0.74), with a negative interaction effect (*ΔCOLD_diet_* l log_2_ FC: –0.81). These results indicate that cold-induced ketone body recycling remains inducible but is dampened under HFD, potentially limiting metabolic flexibility and compensatory lipid synthesis in insulin-resistant BAT. Consistently, only *COLD_HFD_* showed activation of the Type II Diabetes Mellitus signaling pathway under cold stress, reflecting a diabetogenic transcriptional environment marked by insulin resistance and impaired glucose homeostasis [51]. This suggests that cold exposure does not fully overcome metabolic dysfunction in BAT during obesity, potentially limiting its systemic metabolic benefits.

Further, cold-exposed HFD mice uniquely activated Sirtuin signaling, potentially a compensatory response to mitochondrial stress, while suppressing pathways related to mitochondrial dysfunction and white adipose tissue browning, indicating a shift from efficient thermogenesis to cellular maintenance [52]. Further, a selective activation of neuroendocrine pathways (e.g., corticotropin-releasing hormone and orexin signaling) may reflect rewiring of central and peripheral regulatory mechanisms to compensate for reduced intrinsic thermogenic capacity [53]. Concurrent suppression of nitric oxide and neuronal nitric oxide synthase (nNOS) signaling, crucial for vascular perfusion and mitochondrial function, points to impaired BAT structural and vascular adaptation under HFD.

We identified several epigenetic regulators differentially expressed in response to cold and diet. Core factors such as *Tet2*, *Dnmt3*, *Cenpc1*, and *Apobec1* responded similarly under both diets, suggesting a foundational epigenetic program supporting cold-induced chromatin remodeling and transcriptomic flexibility. In contrast to white adipose tissue, where *Mettl4* is upregulated under obesogenic conditions [54], we observed its downregulation in BAT only in *COLD_HFD_*. Given that Mettl4-mediated N6-adenine methylation has been implicated in adipocyte differentiation and lipid production [55], it is conceivable that high-fat diets constrain the differentiation and functional capacity of BAT through epigenetic mechanisms. Conversely, cold exposure decreased *Tet3* and *Dnmt1* expression under chow but not HFD. As Tet3 and Dnmt1 regulate adipocyte expansion and mitochondrial function as well as chromatin architecture, respectively, their cold-induced downregulation may favor thermogenesis by limiting hypertrophy and enhancing mitochondrial and epigenetic adaptability [56–58]. *Dnmt1* deletion in BAT has no metabolic impact under standard diets [59], but may gain relevance under thermogenic or obesogenic stress. A previous study demonstrated the downregulation of *Dnmt1*, *Dnmt3a,* and *Dnmt3b* in BAT upon cold exposure under chow diet conditions, linking reduced DNA methylation to enhanced thermogenic activation [60]. Our findings confirm this pattern under chow conditions but reveal a lack of regulation under HFD, supporting the notion that epigenetic flexibility is impaired in obesogenic environments. *Apobec3*, a cytidine deaminase implicated in RNA and potentially DNA editing [61], was downregulated by cold under both diets, though significantly more under chow. This pattern may reflect its role in facilitating transcriptomic remodeling during thermogenic activation that may be blunted under HFD. Therefore, BAT activation appears to be governed by a conserved core of methylation regulators responsive to cold, while additional, context-specific pathways are modulated by the dietary environment. This dual mechanism supports a model in which a stable epigenetic framework is dynamically adjusted to accommodate metabolic demands, thereby fine-tuning the thermogenic capacity of BAT.

By conducting in vitro methylation analyses, we confirmed the epigenetic transcription control of certain DMEGs and important BAT regulatory genes such as *Bmp8b* or *Cpne5* pointing towards a valid impact of the DNA methylation on BAT function. By using Aza and SAM, we cannot rule out the possibility that other epigenetic mechanisms than DNA methylation led to these observations as especially SAM serves as methyldonor for various processes [62]. In any case, our results strongly support the highly dynamic control of BAT by epigenetic alterations.

### 4.1. Limitations

A limitation arises from age differences between diet groups (11 weeks chow vs. 22 weeks HFD). First, the difference in bodyweight between chow and HFD mice may have partially arisen from the age gap, as there is still a slow increase in body weight beyond adulthood at 12 weeks of age [37]. However, male chow fed mice at 22 weeks of age would reach approximately 30g bodyweight which still is significantly different from mice on HFD at the same age. Second, although BAT function declines with age, mice used in this study are of a comparable age-range related to BAT function. Studies show that mice aged between 12 weeks and six months present a stable metabolic active BAT, with stable mitochondrial content and thermogenic potential before these factors start to decline and lipid accumulation increases [34–36]. Although we cannot bioinformatically address this issue in our comparative analyses, we believe that age is not a main driving factor of the here observed effects as we addressed a comparable age range regarding bodyweight and cold adaptation of BAT. Anyway, regardless of BAT function, global DNA methylation reduces with age while specifically promoter regions tend to get hypermethylated with age [63]. While we cannot address the age factor in our analyses due to group specific age differences, we expect a limited impact of age on our results based on comparing mice in a suitable age range meaning only “young adult mice” [64]. A further limitation is the use of only male mice, which may limit generalizability to females. However, given the known sex-specific differences in adipose biology, BAT activity, and epigenetic regulation, addressing only males is beneficial regarding the addressed methodology. Future studies should include female mice to assess potential sex-by-diet or sex-by-temperature interactions. Small sample sizes (7-8 per group) limit statistical power to detect subtle effects. In addition, RRBS data provides only partial genome coverage, primarily focusing on CpG-rich regions, potentially overlooking important regulatory areas outside these regions. In contrast, whole-genome bisulfite sequencing could provide more comprehensive methylation profiling.

## 5. Conclusions

In conclusion, this study provides the first evidence that cold-induced DNA methylation remodeling in BAT is strongly shaped by dietary context, particularly under obesogenic conditions. Our integrative analysis reveals that, while cold exposure activates a conserved thermogenic transcriptional program, high-fat diet attenuates epigenetic flexibility and restricts full BAT activation. These findings underscore that environmental temperature and dietary intake interact to regulate BAT function via interconnected transcriptional and epigenetic mechanisms. Importantly, we identify both core and context-specific regulators that mediate this response, offering potential molecular targets for future therapies. By demonstrating that obesity alters the epigenetic landscape of cold adaptation, our study opens new avenues for nutritional or epigenetic interventions aimed at restoring BAT plasticity and enhancing thermogenic capacity in metabolic disease.

## Data Availability

Raw RNAseq and RRBS data have been deposited in the Sequence Read Archive (SRA, https://www.ncbi.nlm.nih.gov/sra/) [65] under the BioProject number PRJNA1294891.

## Acknowledgements

We acknowledge the technical support of the Core Unit DNA Technologies of the Faculty of Medicine, University of Leipzig, and thank Dr. Knut Krohn, Stefanie Ziesche and Kathleen Schön. We thank Jenny Schuster and Lisa Gärtner from the animal facility, and Ines Müller, Claudia Gebhardt and Olivia Paetow for excellent technical assistance.

## Author contributions

Conceptualization: TH, AH, KRZ, JTH, and JW; Methodology: AH, TH, KRZ, and JW; Investigation: KRZ, HB, and JW; Validation: KRZ, HB, and JW; Formal analysis: TH; Data curation: TH and AH; Writing – original draft: TH, AH, KRZ, JW, and JTH; Writing – review & editing: TH, AH, KRZ, HB, LM, PK, MS, MB, JTH, and JW; Visualization: TH; Supervision: AH and JTH; Funding acquisition: MB, JW, and JTH.

## Funding

This work was funded or supported by grants of the Deutsche Forschungsgemeinschaft, project number 209933838 (SFB1052 “Obesity Mechanisms”: B1 to MB, C7 to JTH) and by the Formel1 Nachwuchsförderung of the Medical Faculty of the University of Leipzig (JW). HB is supported by a doctoral scholarship of the Studienstiftung des Deutschen Volkes. LM is supported by the Swedish Research Council, the European for the Study of Diabetes and the German Diabetes Association.

## Conflict of interest

MB received honoraria as a consultant and speaker from Amgen, AstraZeneca, Bayer, Boehringer-Ingelheim, Lilly, Novo Nordisk, Novartis, and Sanofi. All other authors declare no conflicts of interest. The funders had no role in the design of the study, the collection, analyses, or interpretation of data, in the decision to publish the results and the writing of the manuscript.

## Abbreviations

Aza: 5’aza-2’-deoxycytidine (Aza)
BAT: Brown adipose tissue
BH: Bonferroni-Holm
DEG: differentially expressed gene
DGE: differential gene expression
DMEG: Differentially methylated and expressed gene
DMP: differentially methylated positions
DMR: differentially methylated regions
eWAT: epididymal white adipose tissue
FC: fold change
FDR: false discovery rate
HFD: high-fat diet
imBA: immortalized brown adipocytes
iWAT: white adipose tissue
PCA: principal component analysis
qPCR: quantitative real-time-PCR
RNAseq: RNA sequencing
RRBS: reduced representation bisulfite sequencing
SAM: S-adenosylmethionine
SVF: stromal vascular fraction
VST: variance stabilization transformation

## Supplementary Figures

**Suppl. Figure 1:**
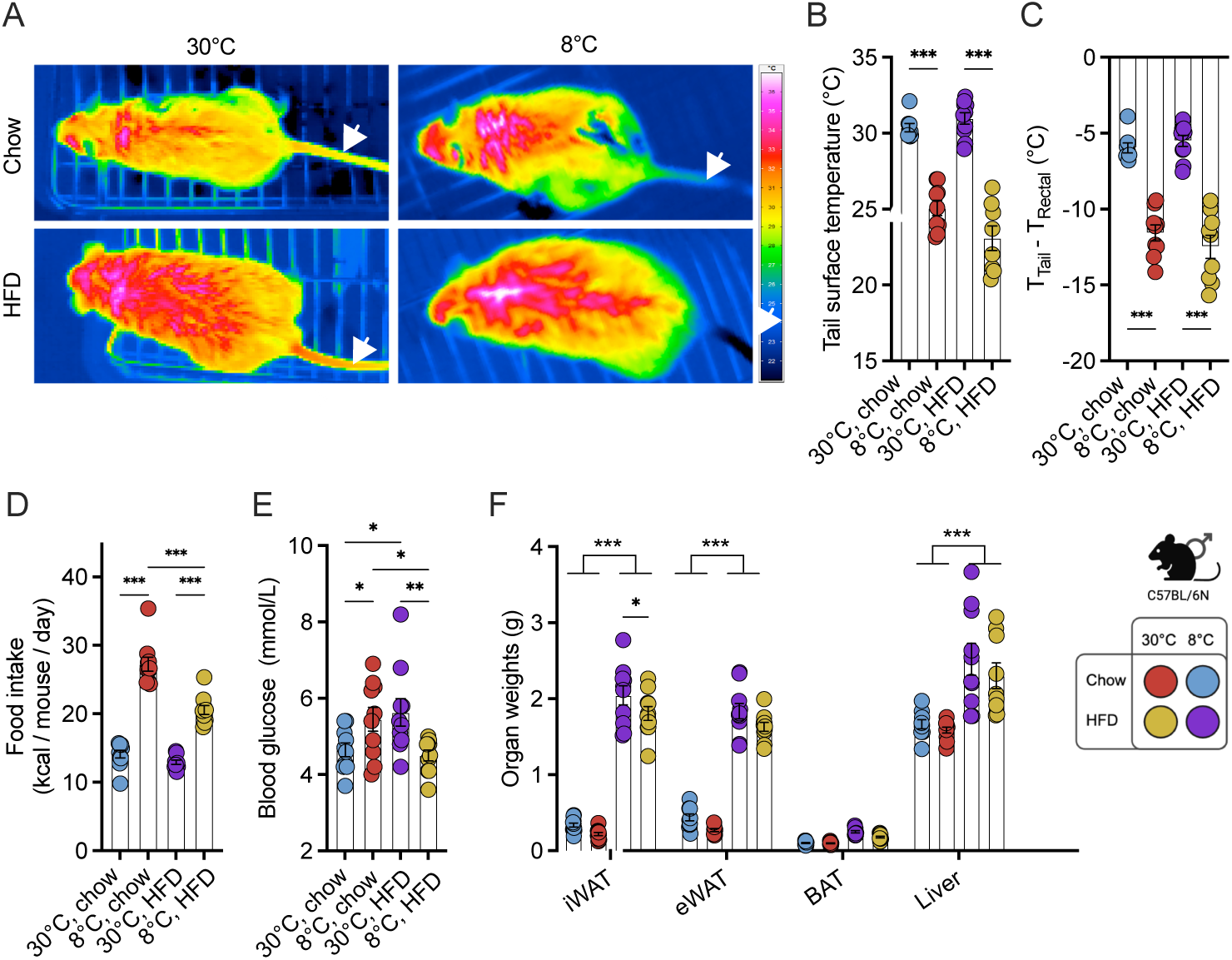
Extended phenotyping of chow and HFD-fed mice housed at 30°C or 8°C. Overview thermal images from brown adipose tissue (BAT) (A), tail surface temperature (B) and heat loss (Ttail – Trectal; (C) in chow and HFD-fed mice housed at thermoneutrality (30°C) or in the cold (8°C) for 7 days. Average daily calory intake (D), final fasted blood glucose (E) and absolute organ weights (F) of inguinal white adipose tissue (iWAT), epididymal white adipose tissue (eWAT), BAT and liver of chow and HFD-fed mice housed at thermoneutrality (30°C) or in the cold (8°C) for 7 days. N = 7-8 per condition. Statistical significance was evaluated one-way ANOVA with Tukey’s post-hoc test (B-D) or uncorrected Fischer’s LDS (E) or two-way ANOVA with uncorrected Fischer’s LDS (F). p <0.05 (*), p <0.01 (**), p <0.001 (***).

**Suppl. Figure 2:**
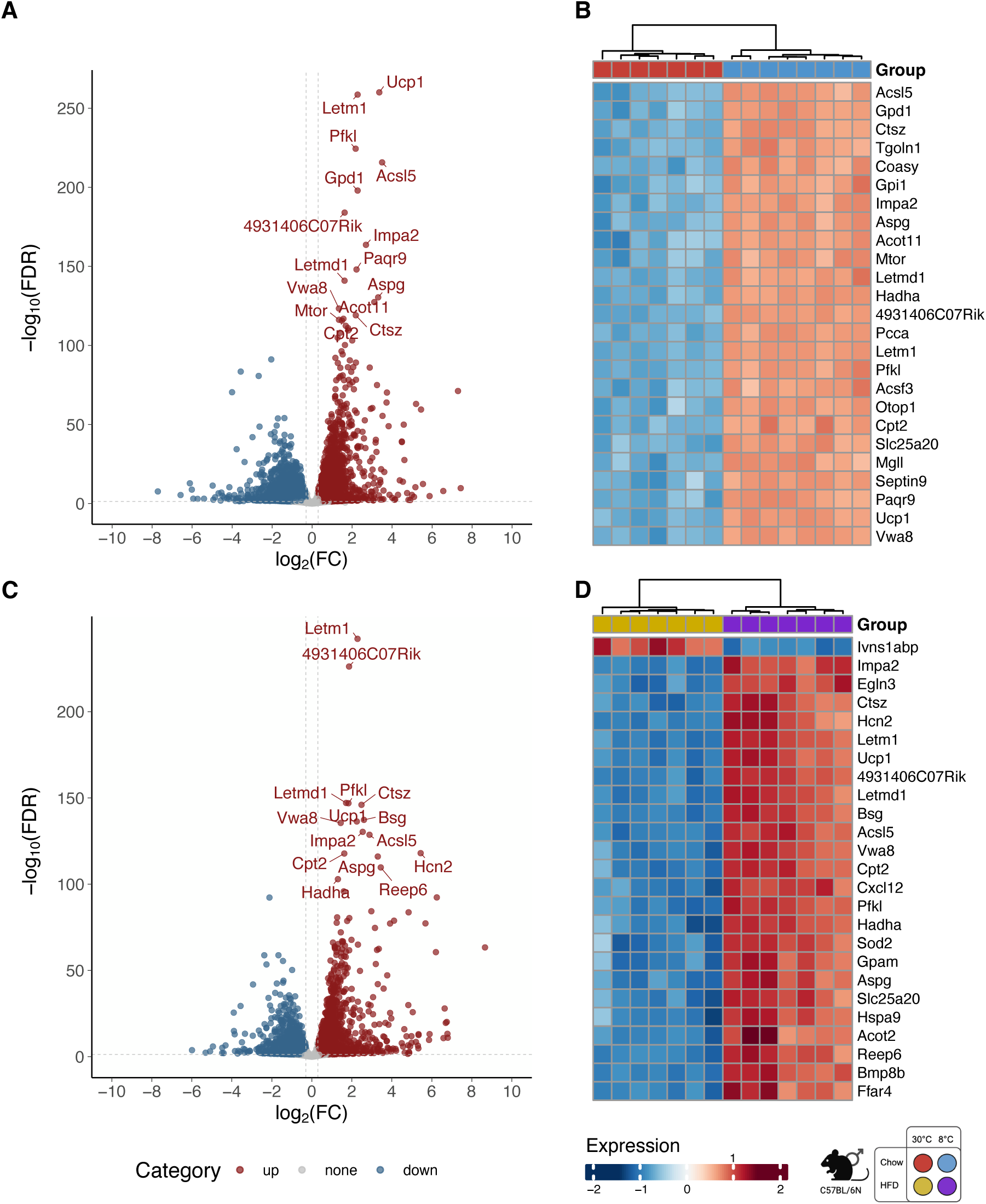
Differential gene expression in response to cold exposure in chow diet and HFD-fed mice. Summary of differential gene expression signals for cold-induced BAT activation in (A, B) normal-weight control mice (*COLD_chow_*) and (C, D) high fat diet (HFD)-induced obese mice (*COLD_HFD_*). Volcano plots (A, C) and heatmaps (B, D) display differentially expressed genes (DGEs) based on log_2_ fold change (FC) and false discovery rate (FDR) signals and show clustering of the top 25 DGEs, respectively.

**Suppl. Figure 3:**
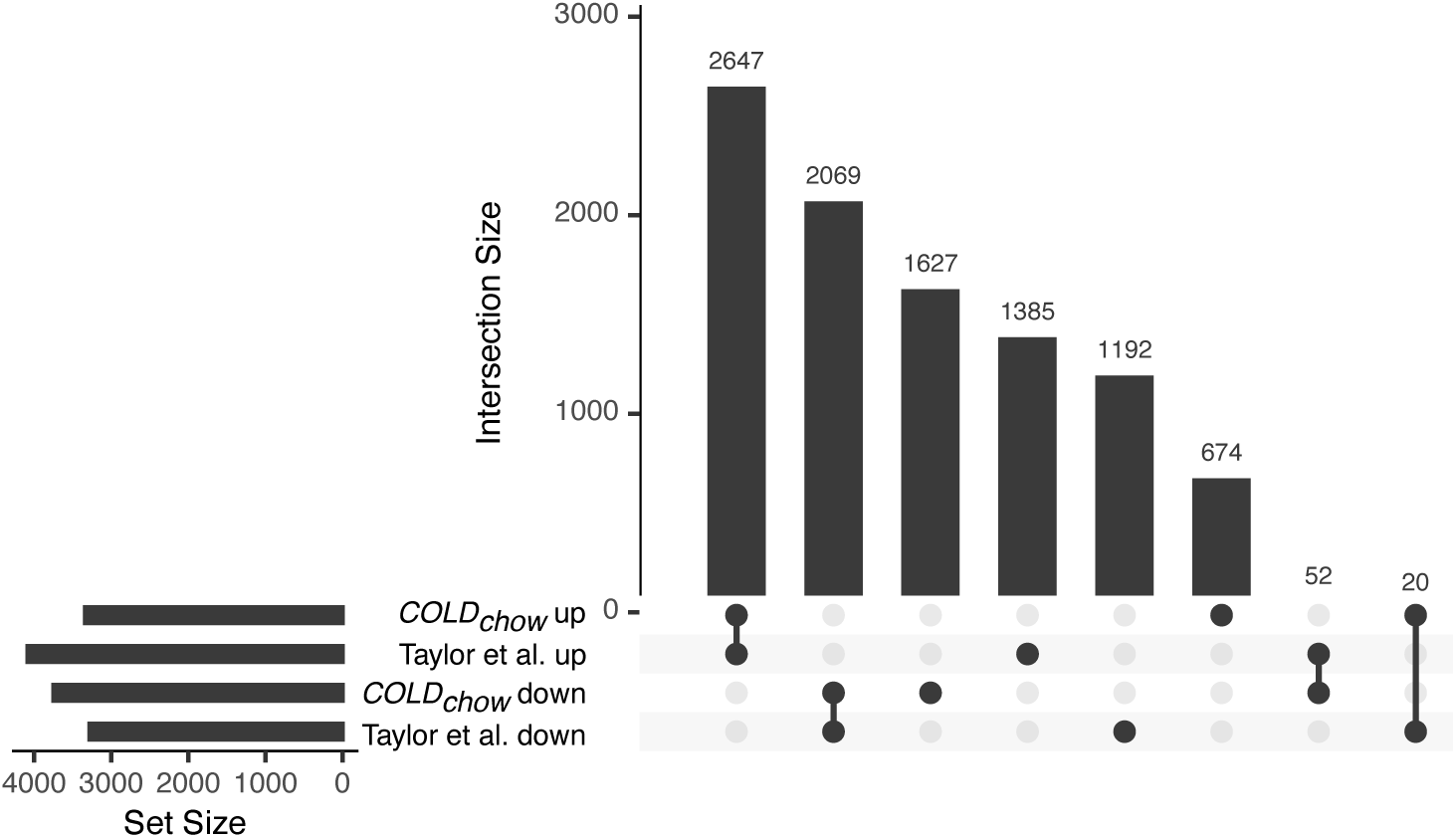
Overlap of DEGs in the *COLD_chow_* condition with published data. Upset plots show the intersection of DEGs from the *COLD_chow_* condition and DEGs reported by Tayler et al. (2024) [60] for chow-fed male mice (8°C vs 28°C), categorized by positive (up) and negative (down) regulation. DEGs with adj. P >0.05 were compared. Set size indicates the total number of DEGs.

**Suppl. Figure 4:**
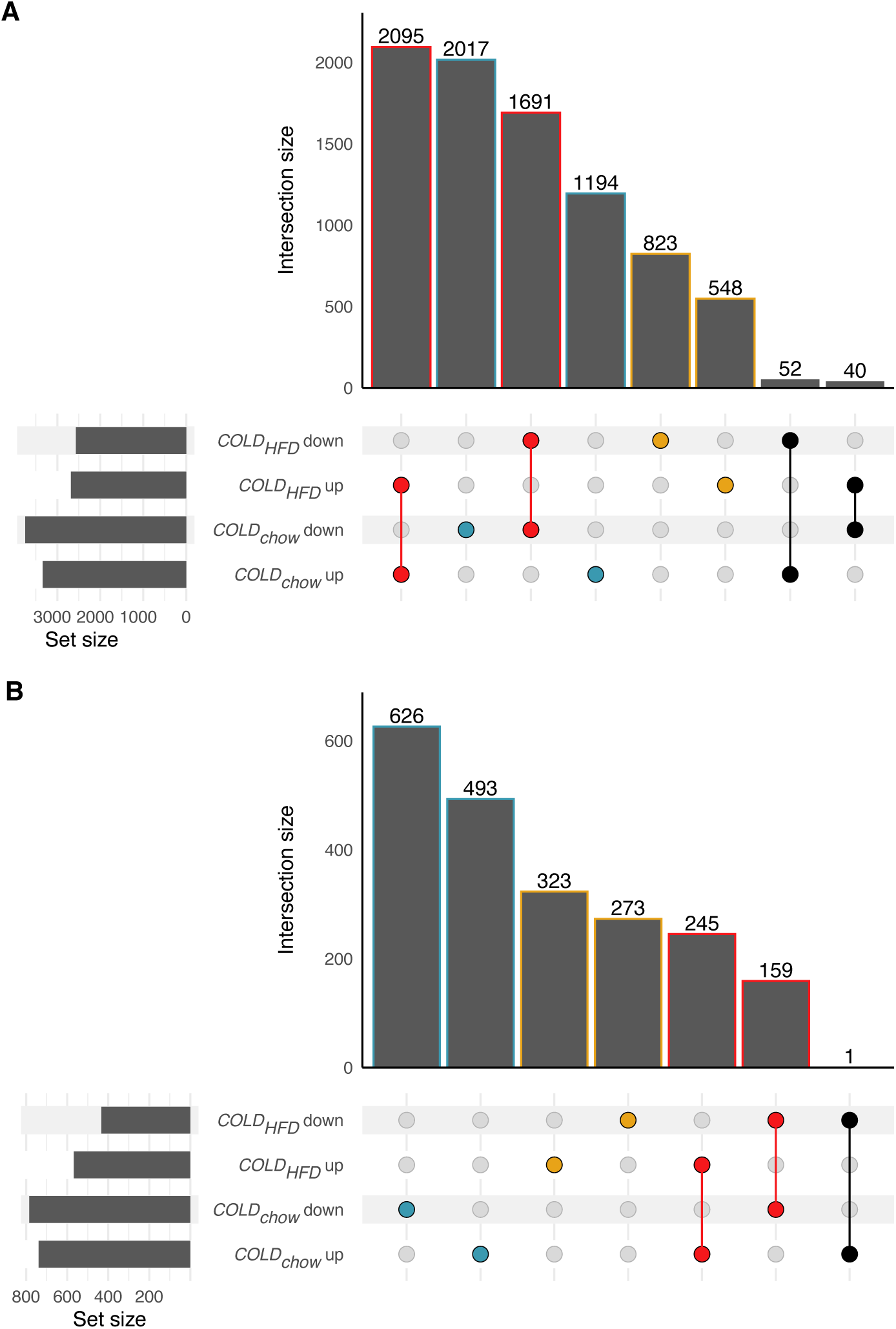
Overlap of DEGs and DMEGs between the *COLD_chow_* and *COLD_HFD_* conditions. Upset plots display the overlap of (A) DEGs and (B) differentially methylated and expressed genes (DMEGs) for the *COLD_chow_* and *COLD_HFD_* condition pairs, categorized by positive (up) and negative (down) regulation. Significant DEGs or DMEGs that show the same and contrary regulatory direction in both conditions are highlighted in red and black respectively. Those significant only in the *COLD_chow_* condition are shown in blue, while those significant only in the *COLD_HFD_* condition are marked in yellow. Genes are counted uniquely, even though multiple DMEGs may be associated with the same gene. Set size indicates the total number of DEGs or DMEGs.

**Suppl. Figure 5:**
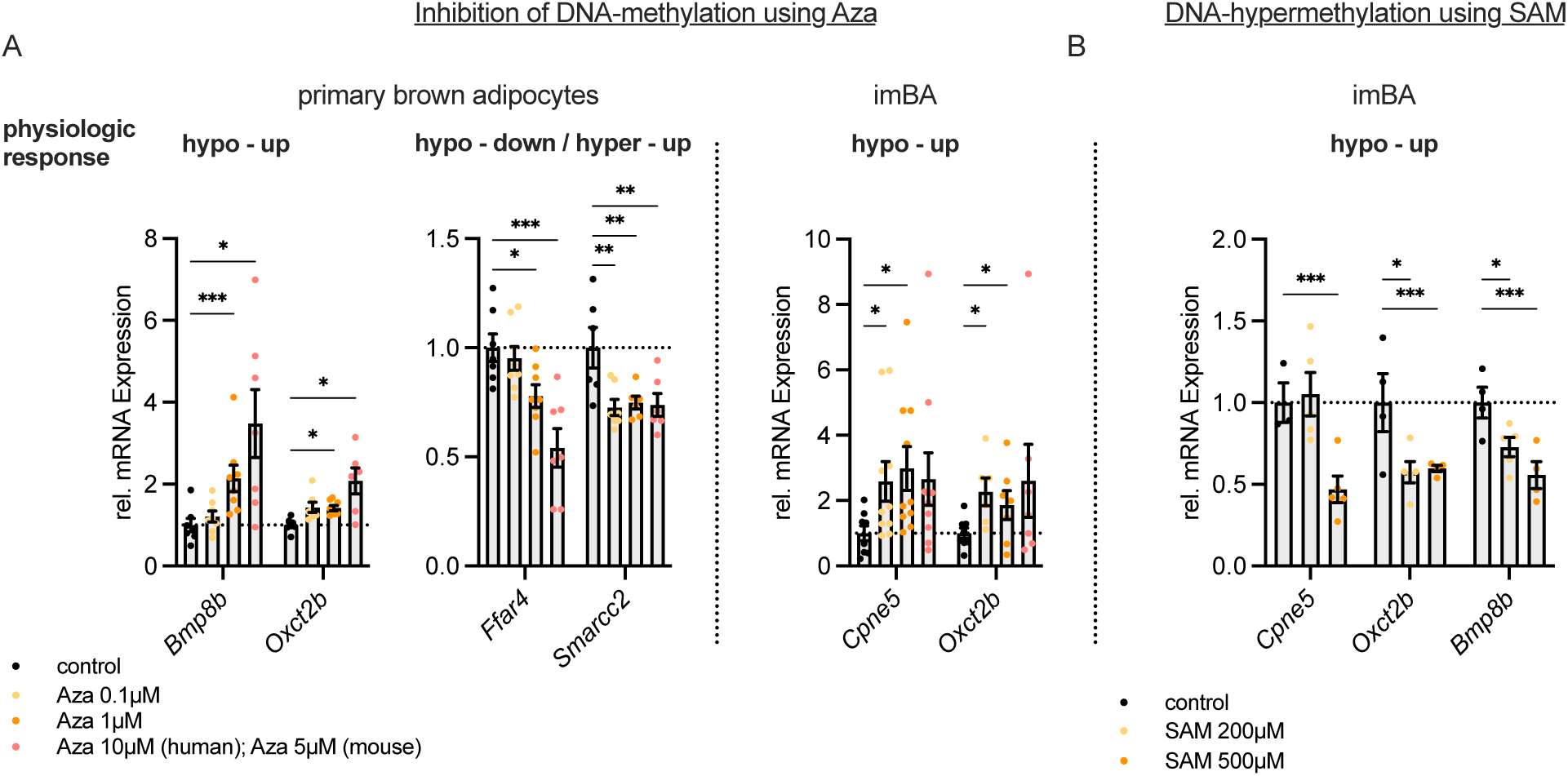
In vitro validation of DNA methylation affecting expression of candidate genes. (A) Inhibition of DNA methylation using indicated concentrations of Aza in mouse immortalized (imBA, left) and primary brown adipocytes (right). Gene expression was analyzed by qPCR, N = 6-10 per condition. (B) Induction of DNA hypermethylation using indicated concentrations of SAM in mouse imBA. Gene expression was analyzed by qPCR, n = 3-5 per condition. Statistical significance was evaluated by two-way ANOVA with uncorrected Fischer’s LSD. p <0.05 (*), p <0.01 (**), p <0.001 (***).

**Suppl. Figure 6:**
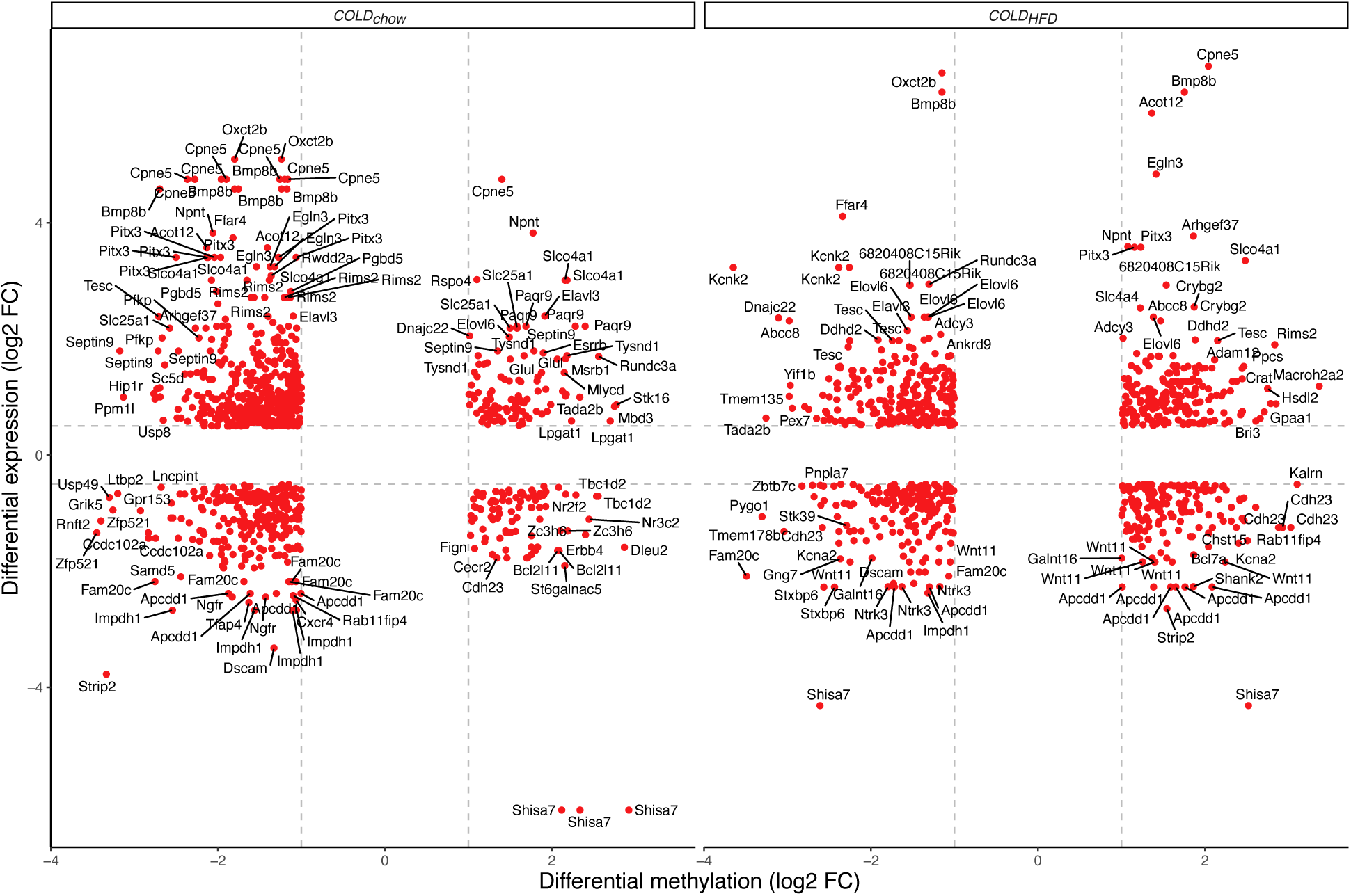
Overlapping DMEGs between *COLD_chow_* and *COLD_HFD_*. The scatter plot displays the log_2_ FC in gene expression and DNA methylation signals for all DMEGs which overlap in both *COLD_chow_* and *COLD_HFD_* condition. Colors correspond to highlights in Suppl. Figure 3. DMEGs with absolute log_2_ FC >0.5 in expression and >1 in methylation signals are shown, while DMEGs in the 5% and 95% quantile of both DMP and DEG log_2_ FC signals are labeled.

**Suppl. Figure 7:**
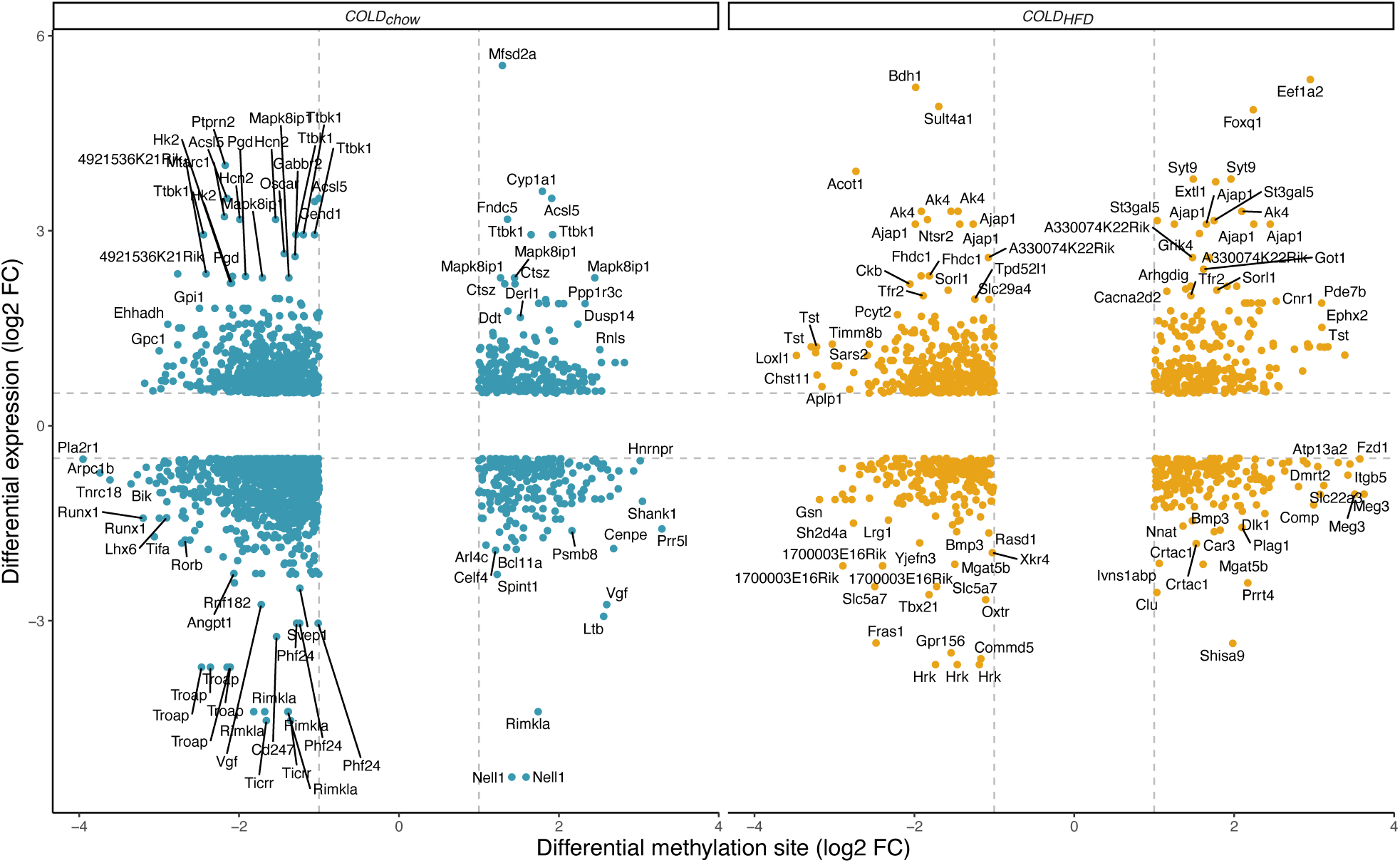
DMEGs that are specifically regulated within each condition. The scatter plot displays the log_2_ FC in gene expression and DNA methylation signals for all DMEGs which are specifically regulated in both *COLD_chow_* and *COLD_HFD_* conditions. Colors correspond to highlights in Suppl. Figure 3. DMEGs with absolute log_2_ FC >0.5 in expression and >1 in methylation signals are shown, while DMEGs in the 5% and 95% quantile of both DMP and DEG log_2_ FC signals are labeled.

## Supplementary Table legends

**Suppl. Table 1:** Overview of primers used for PCR.

**Suppl. Table 2:** Overview of significant (FDR <0.05) DEGs for the *COLD_chow_* contrast. DEGs that overlap with Tayler et al. (2024) [60] (chow-fed mice, 8°C vs 28°C) are listed.

**Suppl. Table 3:** Overview of significant (FDR <0.05) DEGs for the *COLD_HFD_* contrast.

**Suppl. Table 4:** Overview of significant (FDR <0.05) DEGs for the *ΔCOLD_diet_* condition.

**Suppl. Table 5:** Overview of DMEGs in the *COLD_chow_* contrast with absolute expression log_2_ FC >0.5 and absolute methylation log_2_ FC >1, including the cell type assignment derived from single-cell RNAseq data obtained from [38].

**Suppl. Table 6:** Overview of DMEGs in the *COLD_HFD_* contrast with absolute expression log_2_ FC >0.5 and absolute methylation log_2_ FC >1.

**Suppl. Table 7:** Overview of DMEGs in the *ΔCOLD_diet_* contrast with absolute expression log_2_ FC >0.5 and absolute methylation log_2_ FC >1.

**Suppl. Table 8:** Significant IPA enrichment (before p adjustment) for canonical pathways categorized by four groups: those exhibiting similar regulation in both *COLD_HFD_* and *COLD_chow_* condition pairs, and those that are uniquely regulated in each individual condition (*ΔCOLD_diet_*, *COLD_chow_*, and *COLD_HFD_*.)

## Notes

### Summary of Updates

All figures have been formatted and integrated into the manuscript.

